# Neuronal titration of *Snca* via enhancer disruption mitigates disease onset in a Parkinson’s disease mouse model

**DOI:** 10.1101/2025.04.09.648026

**Authors:** Rachel J. Boyd, A Ra Kho, Sarah A. McClymont, Stacie K. Loftus, Han Seok Ko, Andrew S. McCallion

## Abstract

α-synuclein (*SNCA*) is the greatest genetic risk factor for sporadic Parkinson’s Disease (PD). Misfolding and overexpression of SNCA underlie pathognomonic features of PD, including SNCA aggregates and midbrain dopaminergic (mbDA) neurodegeneration. We recently identified an *SNCA* intronic sequence that harbors variation associated with PD risk and demonstrated its role as a neuronal cis-regulatory element (CRE). Here, we engineered a mouse model lacking this sequence, which exhibits significantly reduced *Snca* transcription in mbDA neurons. Employing a battery of motor, molecular, and histological assays in an established mouse model of PD, we demonstrate that mice lacking this *Snca* enhancer are protected against PD-relevant histopathology and motor impairments. By targeting a cell-dependent CRE to diminish PD onset/progression in mice, we introduce a potentially powerful therapeutic avenue for PD.

## INTRODUCTION

Parkinson’s disease (PD) follows Alzheimer’s disease as the second most common age-related neurodegenerative disorder among Americans^1^ and affects over 10 million people worldwide. The clinical presentation of PD typically emerges between 50-60 years of age^2^, whereupon patients generally begin to exhibit progressively debilitating motor deficits, including resting tremor, postural instability, rigidity, and bradykinesia^3^. These symptoms primarily arise due to the preferential loss of midbrain, dopaminergic (mbDA) neurons in the substantia nigra (SN)^3^, in which accumulation of misfolded α-synuclein (SNCA; α-Syn) protein aggregates, called Lewy bodies (LB)^4^, is a pathological hallmark.

The primary function of SNCA is thought to involve modulating neurotransmitter release through effects on the SNARE complex^5,6^. While the precise mechanism of SNCA neurotoxicity in this context is unclear, the effects of *SNCA* overexpression/gene amplification are posited to include loss of presynaptic proteins, decrease in neurotransmitter release, redistribution of SNARE proteins, enlargement of synaptic vesicles, and inhibition of synaptic vesicle recycling^7^. Regardless of the proposed mechanisms, it is well-known that genetic and/or environmental insults^8,9^ can result in *SNCA* structural mutation^10^ or overexpression, both of which promote the formation of intracellular, neurotoxic, protein aggregates that elicit neurodegeneration through oxidative stress and neuroinflammation^7,11,12^.

Furthermore, the *SNCA* locus consistently represents the most significant hit in PD genome-wide association studies (GWASs)^2,13,14^. As with the majority of GWAS-identified functional risk variants, functional variation at this locus is expected to lie within non-coding DNA^15^ and modulate *SNCA* transcriptional control. In line with this model, non-coding variants have already been identified that modulate *SNCA* transcriptional regulatory control, contributing to PD risk^16^.

In a recent study, we identified a region within intron 4 of *SNCA* in which chromatin was preferentially open in mbDA neurons compared to those outside the midbrain^17^. This sequence harbors two, tightly linked variants that are enriched among PD patients, relative to undiagnosed controls, and when tested in transgenic zebrafish and mice, this sequence appeared to function as a putative enhancer that directs expression in catecholaminergic neurons^17^. While PD-associated variants in enhancers of *SNCA* have been identified by us and others^16,17^, the extent to which these PD-associated enhancer sequences modulate PD pathology is understudied.

Here, we assess the biological function of this sequence, as well as its relevance to modulate disease risk. We hypothesize that our identified enhancer sequence is required for transcriptional control of *Snca*, and that it modulates PD risk by modifying *Snca* expression; and thus, the risk of associated inflammatory response. We further theorize that impeding the function of this enhancer will have a protective effect against the progression of PD neuropathology.

To test these hypotheses, we developed a mouse model lacking both copies of this enhancer, as well as littermates that possess one or both intact copies. Mice of all three genotypes were subject to intrastriatal delivery of α-Syn preformed fibrils (PFF), a PD-relevant insult intended to seed PD pathology, or vehicle. Mice were then aged and assayed for the progression of PD-like pathology via established motor and histological assays.

## RESULTS

### Founder mice lacking *Snca*Enh+37 demonstrate reduced *Snca* expression

To test our hypothesis that this neuronal enhancer is required for transcriptional control of *Snca*, we engineered mice lacking this sequence. We designed single guide RNAs (sgRNAs) to flank the *Snca* enhancer interval and induce a 2.76 kb deletion, 37.49 kb downstream from the *Snca* transcription start site within intron 4 (*Snca*Enh+37; **Figure 1A; Table S1; Figure S1**). We identified 5 independent founder mice that were heterozygous for this element, which were progressively backcrossed to C57BL/6J to ensure an isogenic background.

**Figure 1.**
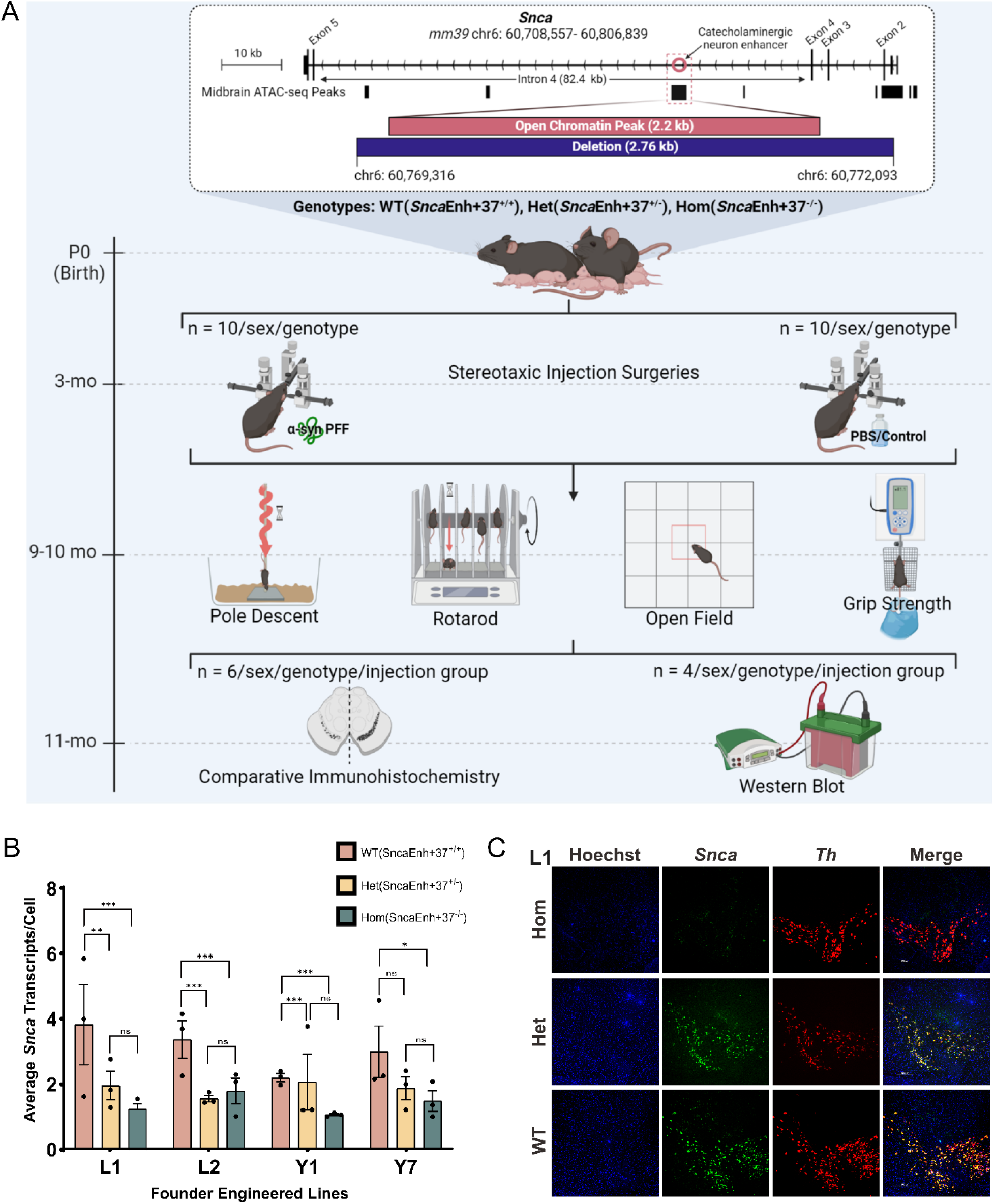
**(A)** Experimental Design. The relative position of the putative enhancer, *Snca*Enh+37, within mouse *Snca* intron 4. This element lies within a differentially accessible chromatin peak (pink) as identified by McClymont *et al*^17^. Mice lacking this element, spanning 2.76 kb (dark blue), were injected with α-syn PFF or PBS control prior to motor, molecular, and histological analyses. **(B)** smFISH/RNAscope reveals a significant reduction in *Snca* transcript among mice lacking *Snca*Enh+37. The bar chart shows the average number of *Snca* transcripts per *Th*+ cell in the SN of 3 mice/genotype/founder line. Error bars represent standard error of the mean (SEM). Significance was determined using a one-way ANOVA and Tukey’s Honestly Significant Difference (HSD) test with a 95% confidence interval. **(C)** Representative smFISH/RNAscope images from *Snca*Enh+37 L1 founder mice. Scale bar = 100μm. See **Table S2** for ANOVA summary statistics. * = p < 0.05, ** = p < 0.01, *** = p < 0.001, ns = not significant.

The degree to which deletion of *Snca*Enh+37 impacts *Snca* expression in mbDA neurons was evaluated via single molecule fluorescent in situ hybridization (smFISH)/RNAscope in four of the founder lines displaying the highest fecundity. We assayed mice across all three genotypes (n = 36; 3/genotype/founder), wild-type /WT(*Snca*Enh+37^+/+^), heterozygous/Het(*Snca*Enh+37^+/-^), and homozygous/Hom(*Snca*Enh+37^-/-^), at N8 generation. These genotypes are designated as WT(*Snca*Enh^+/+^); Het(*Snca*Enh^+/-^), and Hom(*Snca*Enh^-/-^), respectively.

To determine whether our engineered mice displayed a dose-dependent reduction in *Snca* within mbDA neurons, we designed our smFISH/RNAscope assay with probes against *Snca* and tyrosine hydroxylase (*Th*) – the rate-limiting enzyme in dopamine biosynthesis^18^ and a commonly used DA neuron marker. We utilized logic gates to quantify *Snca* transcripts only in DA neurons of the SN, considering neuronal cells that contained DNA (Hoechst+) and were *Th+* (**Figure 1B-C; Figure S2**). We quantified the average number of *Snca* transcripts in these cells, which revealed a consistent reduction in the level of *Snca* transcripts between WT(*Snca*Enh*^+/+^*) and Hom(*Snca*Enh*^-/-^*) mice, and between WT(*Snca*Enh^+/+^) and Het(*Snca*Enh*^+/-^*) mice, thus, confirming the dependence of *Snca* expression in mbDA neurons on *Snca*Enh+37 (**Figure 1B-C; Figure S3; Table S2)**. However, comparison of data from Hom(*Snca*Enh*^-/-^*) and Het(*Snca*Enh*^+/-^*) mice reveal no significant additional reduction upon removal of the second functional enhancer allele (**Figure 1B-C; Figure S3; Table S2**).

Founder line L1 exhibited the largest reduction in *Snca* expression between WT(*Snca*Enh*^+/+^*) and Hom(*Snca*Enh*^-/-^*), as well as a dose-dependent trend of *Snca* reduction between Het(*Snca*Enh*^+/-^*) and Hom(*Snca*Enh*^-/-^*), and was selected for subsequent experimental analysis. Relative to WT(*Snca*Enh*^+/+^*) mice, we observed 2.98 fewer copies of *Snca* transcript per *Th*+ cell in Het(*Snca*Enh*^+/-^*) mice and 3.52 fewer copies of *Snca* per *Th*+ cell in Hom(*Snca*Enh*^-/-^*) mice (*p* < 0.01; **Figure 1B; Table S2**). These data demonstrate that loss of *Snca*Enh+37 is accompanied by a dose-dependent reduction in *Snca* transcription in mbDA neurons; suggesting that sequence within the deleted interval is required for transcriptional modulation of *Snca*.

### *Snca*Enh+37 enhancer deletion mice are protected against PD-relevant motor phenotypes

Next, to test the therapeutic relevance of ablating this sequence, we performed stereotaxic injections surgeries with 10 mice/sex/genotype receiving α-Syn PFF and 10 control mice/sex/genotype receiving saline (PBS) solution, at three months of age (≥N10 backcross generation). α-Syn PFF is an established PD-relevant insult, and previous work has shown that WT(*Snca*Enh*^+/+^*) mice exposed to this insult via intrastriatal injection develop clinically-relevant symptoms of PD, including motor deficits, LB formation, neuroinflammation, and neurodegeneration of DA neurons^19–24^.

Beginning at 6 months post-α-Syn PFF injection, we evaluated n = 10 mice/sex/genotype/injection group with a battery of motor assays, including pole descent, grip strength, and rotarod tests. These are well-established assays for motor impairment and have been widely used to validate the efficacy of α-Syn PFF-induced, PD-like motor phenotypes^20,21,23–27^. In particular, the pole descent test is widely regarded as a sensitive assay of mouse motor deficits^20,21,23–27^.

We hypothesized that PBS control mice would outperform α-Syn PFF-injected mice, regardless of genotype. Further, we predicted that the severity of motor impairment in α-Syn PFF-exposed mice lacking one or both copies of *Snca*Enh+37 would be reduced, relative to α-Syn PFF-exposed mice harboring both intact copies of *Snca*Enh+37. In agreement with previous pole descent data demonstrating that α-Syn PFF injections induce an impaired motor phenotype^20,21,23–26^, we observed induction of PD-like motor phenotypes by pole descent test (**Figure 2A**). WT(*Snca*Enh*^+/+^*) α-Syn PFF mice were >2 s slower to complete the pole descent test than mice in the PBS groups (**Figure 2A**; WT:α-Syn PFF–WT:PBS = 2.08 s, *p* = 0.0385; WT:α-Syn PFF– Het:PBS = 2.34 s, *p* = 0.0113; WT:α-Syn PFF–Hom:PBS = 2.09, *p* = 0.0419). By contrast, Hom(*Snca*Enh*^-/-^*) α-Syn PFF mice and Het(*Snca*Enh*^+/-^*) α-Syn PFF mice performed similarly to PBS injected controls (**Table S3**) and demonstrated an improved pole descent time compared to WT(*Snca*Enh*^+/+^*) α-Syn PFF mice (WT:α-Syn PFF–Het:α-Syn PFF = 2.51 s, *p* = 0.0042; WT:α-Syn PFF–Hom:α-Syn PFF = 2.85 s, *p* = 0.0013). The pole descent data suggests that a lack of *Snca*Enh+37 is more protective in female mice (**Figure S4A; Table S4A**) and does not reach the threshold for statistical significance in male mice (**Table S5)**, despite exhibiting analogous trends (**Figure S4A**). However, this result is likely due to less severe phenotypic induction by α-Syn PFF in male mice. In combination, our pole descent data demonstrates that injection of α-Syn PFF successfully induced motor impairment, as expected, and that mice lacking *Snca*Enh+37 were protected against PD-relevant motor symptom onset.

**Figure 2.**
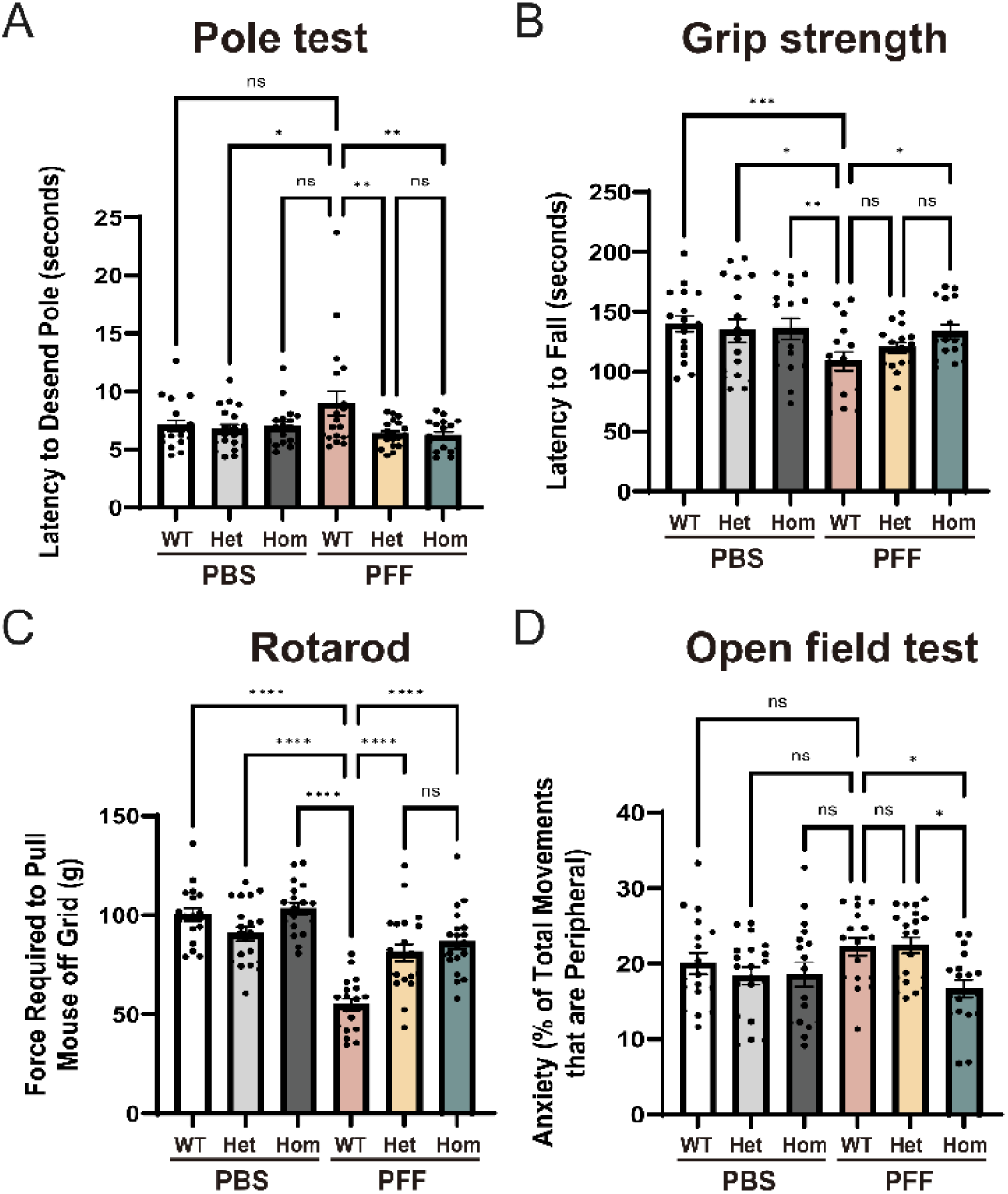
Mice lacking *Snca*Enh+37 are protected against motor deficits as measured by: **(A)** Pole descent performance; **(B)** Grip strength; and **(C)** Rotarod performance. **(D)** Mice lacking *Snca*Enh+37 are also protected against anxiety-like behaviours in the open field test. Error bars represent standard error (SE). Significance was determined using a two-way ANOVA and Tukey’s Honestly Significant Difference (HSD) test with a 95% confidence interval. See **Tables S3-S26** for ANOVA summary statistics. * = p < 0.05, ** = p < 0.01, *** = p < 0.001, **** = p < 0.0001, ns = not significant.

In alignment with previous work showing that α-Syn PFF can induce a motor impairment phenotype measurable by grip strength deficits^20,23,25,26^, α-Syn PFF injected mice were significantly weaker than PBS injected mice (α-Syn PFF-PBS = −23.86, *p* = 1.84E-13; **Table S6**). WT(*SncaEnh^+/+^*) α-Syn PFF mice were the weakest group, requiring 35-49 g less force to pull them from the grid, compared to PBS-treated mice (**Figure 2B**; WT:α-Syn PFF–WT:PBS = −46.13 g, *p* = *p* < 0.0001; WT:α-Syn PFF–Het:PBS = −35.89 g, *p* < 0.0001; WT:α-Syn PFF–Hom:PBS = −48.43 g, *p* < 0.0001). Het(*Snca*Enh*^+/-^*) α-Syn PFF and Hom(*Snca*Enh*^-/-^*) α-Syn PFF mice were significantly stronger than WT(*Snca*Enh*^+/+^*) α-Syn PFF mice (WT:α-Syn PFF–Het:α-Syn PFF = − 26.42 g, *p* = *p* < 0.0001; WT:α-Syn PFF–Hom:α-Syn PFF = −31.65, *p* < 0.0001; **Figure 2B**), consistent with our prediction that enhancer deletion mice will be protected against motor symptom onset and severity. Furthermore, Het(*Snca*Enh*^+/-^*) α-Syn PFF and Hom(*Snca*Enh*^-/-^*) α-Syn PFF mice did not exhibit as severe a reduction in grip strength compared to matched genotype controls (Het:α-Syn PFF-Het:PBS = −9.46 g, *p* = 0.379; Hom:α-Syn PFF-Hom:PBS = −16.78 g, *p* = 0.0129; **Table S6**). These significant trends were observed in both male and female mice (**Tables S7-S8, Figure S4B**), further strengthening the demonstration that *Snca* enhancer deletion mice effectively rescue the parkinsonian motor impairments induced by α-Syn PFF injection.

Intrastriatal α-Syn PFF injections also induced an impaired motor phenotype measurable by rotarod assay^20,21,24,26,27^. PBS control mice were able to remain on the rotarod 16.36 s longer, on average, than their α-Syn PFF injected counterparts (*p* = 0.008; **Figure 2C**; **Table S9-S11**). Specifically, WT(*Snca*Enh*^+/+^*) α-Syn PFF mice demonstrated reduced latency to fall from the rotarod relative to WT(*Snca*Enh*^+/+^*) PBS mice (WT:α-Syn PFF – WT:PBS = −31.165 s; *p* = 0.0404), while mice lacking one or both copies of *Snca*Enh+37 did not exhibit motor impairment, relative to their respective PBS-treated genotype controls (Het:α-Syn PFF – Het:PBS = −15.181 s, *p* = 0.691; Hom:α-Syn PFF – Hom:PBS = −2.389, *p* = 0.9999). However, the data for mice lacking one or both copies of *Snca*Enh+37 trends towards, but falls short of statistical significance in their latency to fall from the rotarod, compared with WT(*Snca*Enh*^+/+^*) α-Syn PFF mice (WT:α-Syn PFF –Het:α-Syn PFF = −10.28 s, *p* = 0.9256; WT:α-Syn PFF –Hom:α-Syn PFF = −24.59 s, *p* = 0.2075; **Figure 2C**). Overall, this rotarod data suggests that our α-Syn PFF injections induced the expected motor impairment in WT(*Snca*Enh*^+/+^*) mice, and that mice lacking SncaEnh+37 were protected against motor symptom onset, relative to their PBS controls.

In summary, WT(*Snca*Enh*^+/+^*) PFF injected mice exhibit onset of PD-like phenotypes, and enhancer deletion mice display marked rescue across almost all evaluated motor phenotypes. Furthermore, Het(*Snca*Enh*^+/-^*) α-Syn PFF and Hom(*Snca*Enh*^-/-^*) α-Syn PFF mice performed similarly to genotype controls in the PBS group on the rotarod, pole descent, and grip strength assays. Taken together, this data suggests that the targeted deletion of *Snca*Enh+37 can protect against the onset and severity of PD-like motor phenotypes in mice, and may represent a potential therapeutic target against PD.

### Mice lacking *Snca*Enh+37 are modestly protected against elevated anxiety-like behaviours

Increased anxiety is associated with clinical progression of PD – and in mice, this may be evaluated in an open field assay in which increased time spent at the periphery of the open field chamber is a sign of elevated anxiety^28^. WT(*Snca*Enh*^+/+^*) mice recorded significantly higher anxiety scores than Hom(*Snca*Enh*^-/-^*) mice, spending 3.5% more time at the periphery of the open field chamber (*p* = 0.0191; **Tables S12-S14**). Hom(*Snca*Enh*^-/-^*) α-Syn PFF mice exhibit a reduction in anxiety-like behaviours. Specifically, Hom(*Snca*Enh*^-/-^*) α-Syn PFF mice spent less time at the periphery of the chamber than WT(*Snca*Enh*^+/+^*) α-Syn PFF mice (WT:α-Syn PFF – Hom:α-Syn PFF = 5.56%, *p* = 0.0309) and Het(*Snca*Enh*^+/-^*) α-Syn PFF (Hom:α-Syn PFF – Het:α-Syn PFF = −5.74%, *p* = 0.0183) mice (**Figure 2D; Tables S12-S14)**. Further, Hom(*Snca*Enh*^-/-^*) PFF mice exhibit similar anxiety levels to PBS controls (Hom:α-Syn PFF–WT:PBS = −3.34%, *p* = 0.4406; Hom:α-Syn PFF–Het:PBS = −1.70%, *p* = 0.9285; Hom:α-Syn PFF–Hom:PBS = −1.88%, *p* = 0.9040); however, WT(*Snca*Enh*^+/+^*) α-Syn PFF and Het(*Snca*Enh*^+/-^*) α-Syn PFF mice also did not exhibit significant differences from PBS controls (WT:α-Syn PFF–WT:PBS = 2.23%, *p* = 0.8201; Het:α-Syn PFF–Het:PBS = 4.04%, *p* = 0.1805). Although *Snca*Enh+37 deletion may protect against the onset and severity of anxiety, the lack of a more pronounced phenotypic induction in WT(*Snca*Enh*^+/+^*) and Het(*Snca*Enh*^+/-^*) α-Syn PFF mice render this observation inconclusive at this stage. α-Syn PFF injected animals recorded more peripheral and, as a consequence, more total movement (locomotion) in the open field chamber compared to PBS mice (*p* < 0.05; **Figure S5-S6; Tables S15-S20**); however, there was no significant impact of α-Syn PFF/PBS injection, genotype, or their interactive effects, on center movements or rearing behaviours (**Figures S7-S8; Table S21-S26**).

### Mice lacking *Snca*Enh+37 are protected against histopathological hallmarks of PD

Lewy Body (LB) formation and DA neurodegeneration are pathognomonic features for PD. Hyperphosphorylation of the SNCA protein, specifically on amino acid residue serine-129 (S129), alters the solubility properties of SNCA; thus, promoting protein aggregation, abnormal function, and neurotoxicity^29^. Phosphorylated S129 (pS129) is a marker for these protein aggregates, which are the primary constituent of LBs. Therefore, to test whether targeted deletion of *Snca*Enh+37 is neuroprotective against LB formation and DA neuron neurodegeneration, we quantified pS129 and Th+ neurons in the midbrains of 6 mice/sex/genotype/treatment using immunohistochemistry (IHC), unbiased stereology, and western blot (WB).

α-Syn PFF-treated mice lacking *Snca*Enh+37 were protected against LB formation and neurodegeneration, compared to the induction of these pathological hallmarks in WT(*Snca*Enh*^+/+^*) α-Syn PFF mice. We performed IHC and quantified LB-like, pS129+ inclusions in Th+ DA neurons **(Figure 3A-C**). We observed a significant dose-dependent reduction in the number of LB-like, pS129+ inclusions in Het(*Snca*Enh*^+/-^*) α-Syn PFF and Hom(*Snca*Enh*^-/-^*) α-Syn PFF mice (WT:α-Syn PFF – Het:α-Syn PFF = 6.458, *p* <0.0001; WT:α-Syn PFF – Hom:α-Syn PFF = 9.758, *p* <0.0001) (**Figure 3B; Table S27**), which corresponded with a dose-dependent increase in Th+ DA neurons (WT:α-Syn PFF – Het:α-Syn PFF = −4.502, *p* <0.0001; WT:α-Syn PFF – Hom:α-Syn PFF = −11.62, *p* <0.0001) **(Figure 3C; Table S28)**. This result represents a near complete rescue of the proportion of Th+ DA neurons that contain LB-like, pS129 inclusions in Het(*Snca*Enh*^+/-^*) α-Syn PFF (WT:α-Syn PFF – Het:α-Syn PFF = 2.403, *p* <0.0001, 76.8% reduction) and Hom(*Snca*Enh*^-/-^*) α-Syn PFF mice (WT:α-Syn PFF – Hom:α-Syn PFF = 2.934, *p* <0.0001, 93.8% reduction), relative to WT(*Snca*Enh*^+/+^*) α-Syn PFF mice (**Figure 3D; Table S29**).

**Figure 3.**
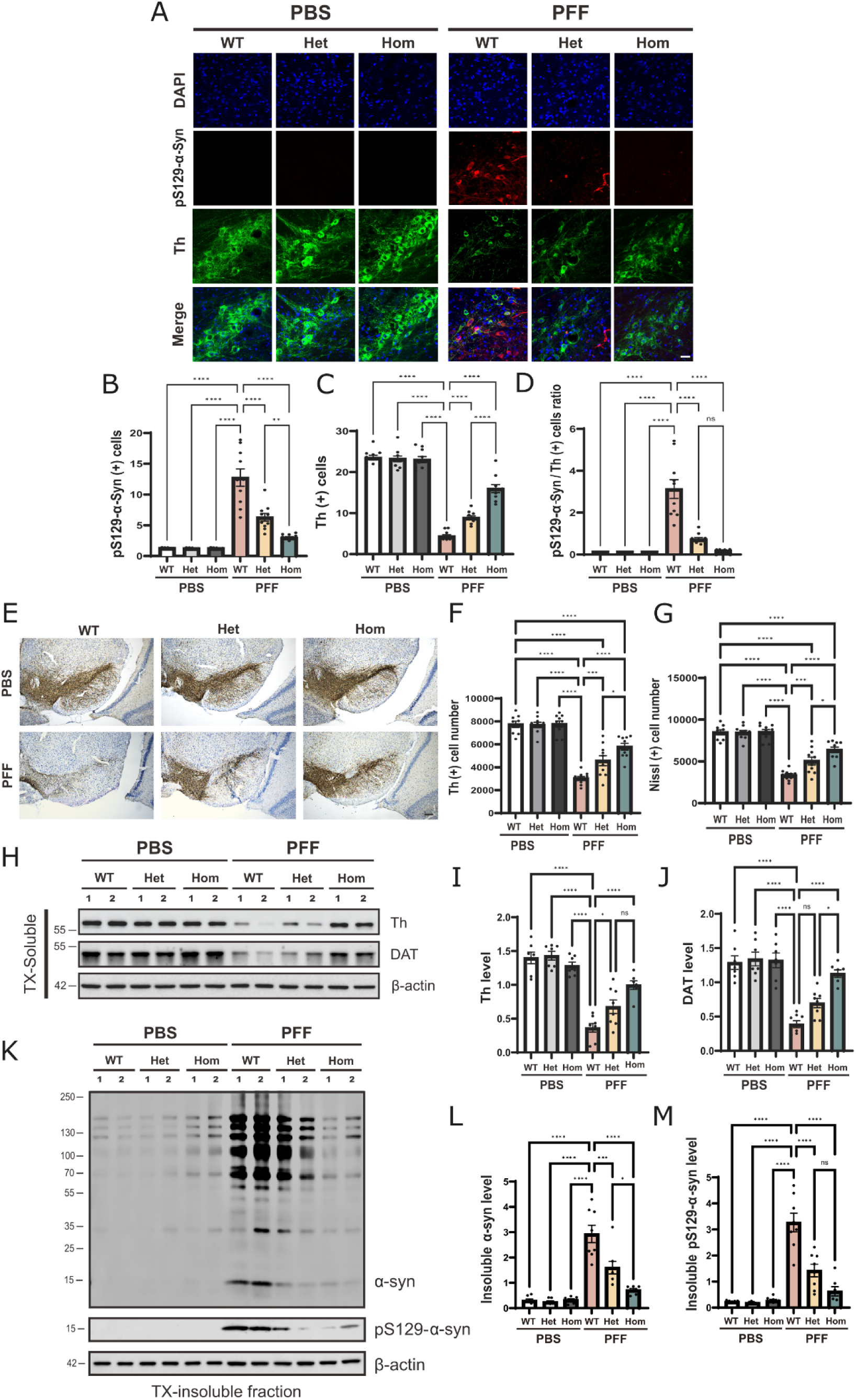
Mice lacking *Snca*Enh+37 are protected against PD-like histopathological phenotypes: **(A)** Representative immunostaining for α-synuclein phosphorylated serine 129 (pS129-α-syn), indicative of LB inclusions, and Th+ dopaminergic neurons in the SN. Scale bar=20µm **(B)** Quantification of pS129-α-syn + immunostained neurons in the SN. **(C)** Quantification of Th+ immunostained neurons in the SN. **(D)** Proportion of Th+ neurons that contain pS129-α-syn/LB inclusions in the SN. **(E)** Representative photomicrographs of coronal sections showing Th+ and Nissl+ neurons in the SN. Scale bar=40µm. **(F)** Unbiased stereological quantification of Th+ DA neurons in the SN. **(G)** Unbiased stereological quantification of Nissl+ neurons in the SN. **(H)** Western blot of Th and DAT from TX-soluble fraction. **(I)** Relative quantification of Th, normalized to β-actin. **(J)** Relative quantification of DAT, normalized to β-actin. **(K)** Western blot from TX-insoluble fraction. **(L)** Relative quantification of insoluble α-syn, normalized to β-actin. **(M)** Relative quantification of insoluble pS129-α-syn, normalized to β-actin. Significance was determined using a two-way ANOVA and Tukey’s Honestly Significant Difference (HSD) test with a 95% confidence interval. See **Tables S27-31** (IHC); **S32-S35** (WB) for ANOVA summary statistics. * = p < 0.05, ** = p < 0.01, *** = p < 0.001, **** = p < 0.0001, ns = not significant.

As expected, WT(*Snca*Enh*^+/+^*) α-Syn PFF mice exhibited a significant loss of midbrain DA neurons; however, in Het(*Snca*Enh*^+/-^*) and Hom(*Snca*Enh*^-/-^*) α-Syn PFF mice, DA neuron loss was significantly reduced (**Figure 3A,C; Table S28**). To further validate these findings, we performed unbiased stereological counting of Th+ neurons in the SN, combined with IHC analysis (**Figure 3E-G)**. Consistent with prior findings^19,27^, WT(*Snca*Enh*^+/+^*) α-Syn PFF mice exhibited significant loss of Th+ (WT:PBS – WT:PFF = 4685, *p* <0.0001; **Table S30**) and Nissl+ (WT:PBS – WT:PFF = 5108, *p* <0.0001; **Table S31**) neurons in the SN, whereas Het(*Snca*Enh*^+/-^*) and Hom(*Snca*Enh*^-/-^*) α-Syn PFF mice possessed significantly more remaining Th+ (WT:PFF – Het:PFF = −1547, *p* = 0.0009; WT:PFF – Hom:PFF = −2761, *p* <0.0001; **Table S30**) and Nissl+ (WT:PFF – Het:PFF = −1712, *p* = 0.0008; WT:PFF – Hom:PFF = −3047, *p* <0.0001; **Table S31**) neurons in the SN, compared to WT(*Snca*Enh*^+/+^*) α-Syn PFF mice (**Figure 3E-G)**.

Next, we conducted WB to validate the levels of Th and dopamine transporter (Slc6a3/Dat1; DAT) in tissue from the ventral midbrain of these mice. Consistent with our IHC results, we observed a significant increase in Th protein levels in Het(*Snca*Enh*^+/-^*) and Hom(*Snca*Enh*^-/-^*) α-Syn PFF mice, compared to WT(*Snca*Enh*^+/+^*) α-Syn PFF mice (**Figure 3H-I**; WT:PFF – Het:PFF = −0.3125, *p* = 0.0408; WT:PFF – Hom:PFF = −0.6319, *p* <0.0001; **Table S32**); corresponding with a comparable increase in DAT levels in Het(*Snca*Enh*^+/-^*) and Hom(*Snca*Enh*^-/-^*) α-Syn PFF mice, compared to WT(*Snca*Enh*^+/+^*) α-Syn PFF mice (**Figure 3H,J**; WT:PFF – Het:PFF = −0.3083, *p* = 0.1134; WT:PFF – Hom:PFF = −0.7584, *p* <0.0001; **Table S33**). Taken together, these results suggest that mice lacking *Snca*Enh+37 are protected against PD-relevant DA neurodegeneration (**Figure 3H-J**).

To further investigate the effect of lacking *Snca*Enh+37 on LB-like pathology, the midbrain of α-Syn PFF-injected mice was fractioned to yield the TX-insoluble fraction, followed by WB analysis. As expected, insoluble α-Syn aggregates and pS129-α-Syn were nearly undetectable in PBS-treated animals, but highly detectable in WT(*Snca*Enh*^+/+^*) α-Syn PFF mice (**Figure 3K**). Compared to WT(*Snca*Enh*^+/+^*) α-Syn PFF mice, TX-insoluble α-Syn aggregates were significantly reduced in Het(*Snca*Enh*^+/-^*) and Hom(*Snca*Enh*^-/-^*) α-Syn PFF mice, in a dose-dependent manner (WT:PFF –Het:PFF = 1.332, *p* = 0.0108; WT:PFF – Hom:PFF = 2.313, *p* = 0.0002; **Table S34; Figure 3K-M**). Specifically, insoluble pS129-α-Syn levels were significantly reduced by 56.4% in Het(*Snca*Enh*^+/-^*) and 80.9% in Hom(*Snca*Enh*^-/-^*) α-Syn PFF mice, compared to the WT(*Snca*Enh*^+/+^*) α-Syn PFF mice (WT:PFF – Het:PFF = 1.843, *p* = 0.0011; WT:PFF – Hom:PFF = 2.728, *p* <0.0001; **Table S35**; **Figure 3K-M)**. Collectively, these results demonstrate that mice lacking *Snca*Enh+37 are protected against neurotoxic α-Syn aggregation and the formation of LB-like inclusions.

### Mice lacking SncaEnh+37 are protected against neuroinflammatory phenotypes

Neuroinflammation is another pathological hallmark of PD and other neurodegenerative disorders^12^. Therefore, to test whether the targeted deletion of *Snca*Enh+37 protects against neuroinflammation in our PD model, we performed IHC and WB analyses against the two major neuroinflammatory cell types, astrocytes and microglia. Both microglia (Iba1) and astrocyte (Gfap) markers showed nearly absent immunoreactivity in all PBS-treated mice (**Figure 4A-C**). By contrast, Iba1 (WT:PBS – WT:PFF = −15.05, *p* <0.0001, **Table S36)** and Gfap (WT:PBS – WT:PFF = −16.18, *p* <0.0001, **Table S37**) were both significantly increased in the SN of WT(*Snca*Enh*^+/+^*) α-Syn PFF mice. Compared to WT(*Snca*Enh*^+/+^*) α-Syn PFF mice, Iba1 (**Figure 4A-B**; WT:PFF – Het:PFF = 8.215, *p* < 0.0001; WT:PFF – Hom:PFF = 11.4, *p* < 0.0001; **Table S36**) and Gfap (**Figure 4A,C**; WT:PFF – Het:PFF = 10.47, *p* < 0.0001; WT:PFF – Hom:PFF = 13.96, *p* < 0.0001; **Table S37**) were significantly reduced in the SN of Het(*Snca*Enh*^+/-^*) α-Syn PFF and Hom(*Snca*Enh*^-/-^*) α-Syn PFF mice, in a dose-dependent manner, as assessed by IHC analysis (**Figure 4A-C**).

**Figure 4.**
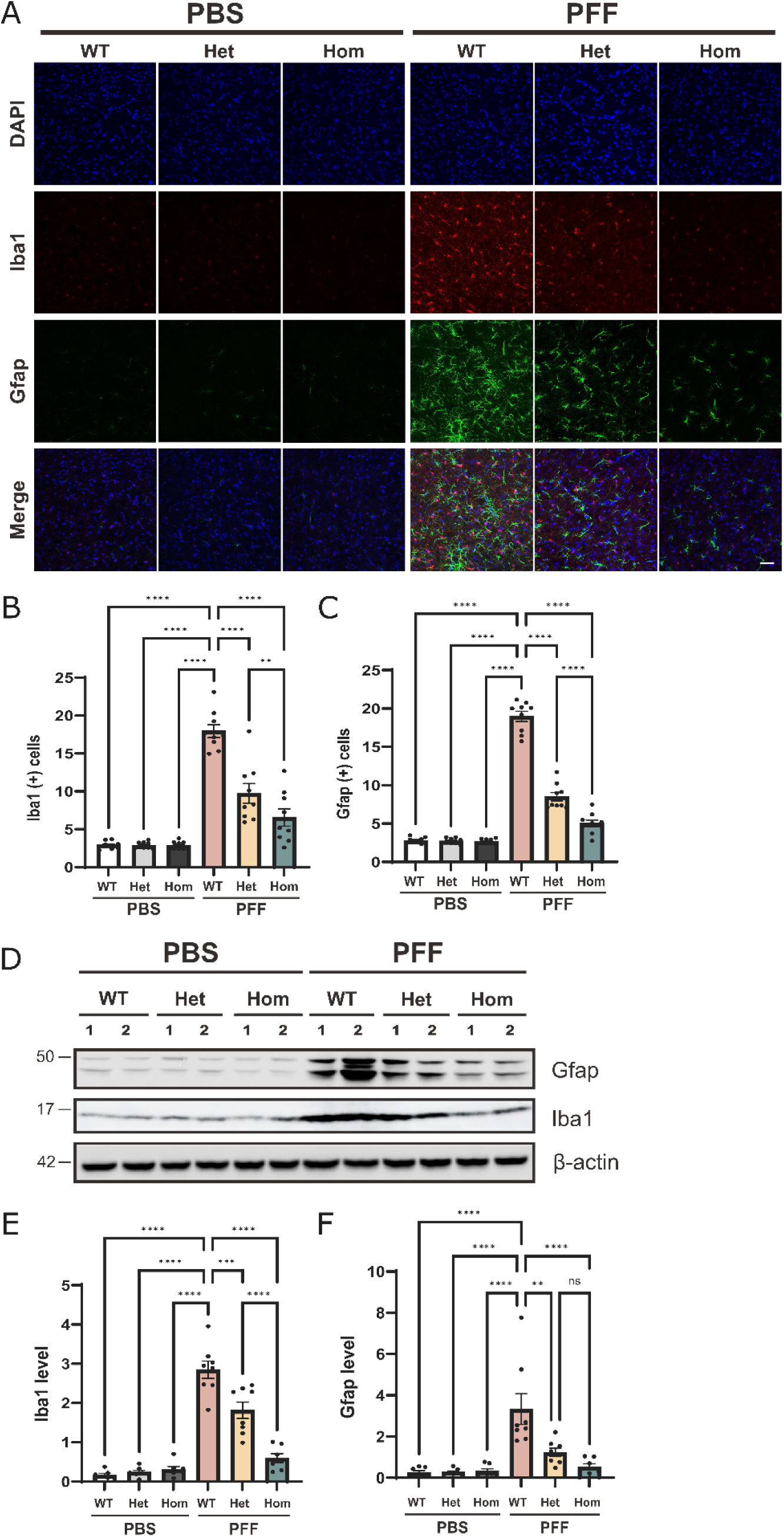
Mice lacking *Snca*Enh+37 are protected against neuroinflammatory phenotypes: **(A)** Representative immunostaining of Iba1 and Gfap in the SN. Scale bar=40µm. **(B)** Quantification of Iba1+ immunostained cells. **(C)** Quantification of Gfap+ immunostained cells. **(D)** Western blot of neuroinflammatory markers, Iba1 (microglia) and Gfap (astrocytes) in the SN **(E)** Relative quantification of Iba1, normalized to β-actin. **(F)** Relative quantification of Gfap, normalized to β-actin. Significance was determined using a two-way ANOVA and Tukey’s Honestly Significant Difference (HSD) test with a 95% confidence interval. See **Tables S36-S37** (IHC); **S38-S39** (WB) for ANOVA summary statistics * = *p* < 0.05, ** = *p* < 0.01, *** = *p* < 0.001, **** = *p* < 0.0001, ns = not significant.

Consistent with our IHC observations, WB analysis revealed nearly absent Iba1 and Gfap protein levels in the MB of PBS treated mice (**Figure 4D-F**), significantly elevated Iba (WT:PBS – WT:PFF = −2.738, *p* <0.0001, **Table S38)** and Gfap (WT:PBS – WT:PFF = −3.205, *p* <0.0001, **Table S39)** levels in the midbrain of WT(*Snca*Enh*^+/+^*) α-Syn PFF mice, and significantly diminished Iba (WT:PFF – Het:PFF = 1.034, *p* = 0.0003; WT:PFF – Hom:PFF = 2.310, *p* <0.0001; **Table S38**) and Gfap (WT:PFF – Het:PFF = 2.100, *p* = 0.0020; WT:PFF – Hom:PFF = 2.936, *p* < 0.0001; **Table S39**) in Het(*Snca*Enh*^+/-^*) α-Syn PFF and Hom(*Snca*Enh*^-/-^*) α-Syn PFF mice. These assays confirm that modulation of *Snca* expression in mbDA neurons by disruption of *Snca*Enh+37 significantly reduces clinically relevant, neuroinflammatory pathology in a mouse model of PD.

## DISCUSSION

We demonstrate that a cis-regulatory element (CRE)/enhancer, with previously identified PD-associated risk variants therein^17^, is a transcriptional regulator of *Snca*. Using a mouse model of PD, we demonstrate that the deletion of this neuron-dependent CRE elicits significant protection from α-Syn aggregation, significant protection from neuroinflammation and neuronal loss, and significantly improved motor function compared to mice in which the CRE remains intact. By disrupting this CRE, we are able to titrate cognate gene transcription, thereby reducing the death of DA neurons and diminishing markers of neuroinflammatory microglia and astrocytes to near-normal levels. This observation coincides with a reduction in both glial cell activation and associated deleterious effects that lead to decreased progression of neurodegenerative symptoms. As such, this novel approach offers a potentially powerful and novel therapeutic avenue that has the potential to extend to synucleinopathies at large, including PD, multiple system atrophy, and dementia with Lewy bodies.

This strategy builds on a foundation of data demonstrating that *SNCA* overexpression promotes misfolding and aggregation in driving PD risk^7,10,17^. Similarly, the requirement for baseline levels of WT *Snca* in mouse models has also been established^20^. Here, we leverage evidence from GWAS, predicting a pathogenic role for regulatory variation at *SNCA*^2,13,14^. We identified a key CRE target at *SNCA/Snca* for therapeutic manipulation, proposing that protective levels of *Snca* modulation might be achieved via ablation of a cell-dependent regulatory sequence at this critical gene.

This critical proof-of-concept study establishes that prophylactic targeted modulation of a CRE effectively ameliorates clinical onset and severity of PD symptoms. We are actively engaged in studies to evaluate the application of this approach to therapeutic windows for intervention after onset of clinical symptoms, and we recognize that, at this stage, we cannot yet address questions of therapeutic delivery. Similarly, we recognize that the potential power of this approach demands continued verification across a spectrum of PD-relevant insults, both genetic and environmental.

Additionally, PD often presents with a variety of non-motor phenotypes, including anosmia, constipation, depression, and REM sleep disorders, which have been shown to precede motor phenotypes by up to 20 years^30^. This prodromal phase of PD suggests that neurodegeneration begins long before the onset of pathognomonic symptoms. In fact, by the time these motor symptoms arise and a clinical diagnosis is made, up to 60-80% of mbDA neurons in the SNpc have already been lost^3^. As such, it is imperative that these findings, and the identification of clinical biomarkers, are fully integrated in the identification of available treatment windows. Understanding how and when key PD genes are regulated will provide insight into the mechanisms of PD risk and progression and can inform the development of broadly effective therapeutic strategies. Further, the extent of protection against non-motor PD co-morbidities (cognitive decline, sleep disruption, *etc.*) provided by this strategy are a matter of ongoing study.

## CONCLUSION

In summary, we show that CRISPR-mediated ablation of a neuronal *Snca* enhancer provides protection against motor deficits and neurodegeneration in a well-established PD mouse model, likely by reducing baseline levels of *Snca* transcription in the midbrain. We leverage gene editing to modulate regulatory element function and in turn titrate the levels of a key gene in PD risk and progression, in a cell-dependent manner to ameliorate the clinical and molecular manifestations of a PD mouse model. The success of this novel strategy has broad implications for the future of therapeutic interventions for PD, other synucleinopathies, and neurodegenerative disorders, and opens the door to a new generation of therapeutic modalities for rare and common diseases alike.

## AUTHOR CONTRIBUTIONS

A.S.M., R.J.B., and H.S.K. conceptualized the study. S.K.L., S.A.M., and A.S.M. designed the mouse model. S.K.L. generated the mouse model. R.J.B., A.S.M., and S.A.M. maintained the mouse model. R.J.B. and A.S.M. backcrossed the mouse model, and R.J.B. validated the resulting inbred lines by PCR and smFISH/RNAscope. A.R.K. undertook quality control of α-Syn PFF and established its cytotoxicity prior to performing α-Syn PFF/PBS injection surgeries. R.J.B. and A.R.K. performed all motor and histological assays. R.J.B. and A.R.K. performed bioinformatics and statistical analyses. R.J.B. wrote and assembled the manuscript, which was revised with input from all authors. R.J.B, A.R.K., H.S.K., and A.S.M. were responsible for the scientific discussion, data interpretation, and iterative revision of the paper. All authors have read and agreed to the published version of the manuscript.

## Supporting information

RNAScope quantification

## ACKNOWLEDGEMENTS

The authors would like to acknowledge Sang Ho Kwon for technical support with RNAscope analysis; Dr. Chantelle Terrillion, the director of the Behavioral Core of the School of Medicine, for assistance with mouse phenotyping; and the Johns Hopkins Research Animal Resources technicians and veterinarians for assistance with mouse husbandry. We also thank Dr. Lauren Jantzie, Dr. Hawley Helmbrecht, and Nelson Barrientos for helpful discussions in the assembly of this work. Figures were created using Biorender.com, R (v4.3.3), and GraphPad Prism (v10.3.0).

## MATERIALS & METHODS

### Generation of Snca Enhancer Deletion Mouse Model

All mice were maintained on a 12-hour light-dark cycle in a temperature and humidity-controlled facility with *ad libitum* access to food and water. All experiments were performed in strict accordance with protocols approved by the Institutional Animal Care and Use Committee (IACUC) at the Johns Hopkins University School of Medicine (MO18M427; MO23M44; MO24M315), which were developed in accordance with National Research Council guidelines.

*Snca* enhancer deletion founder mice (C57BL/6J-*Snca^em1Asm^*/J, or *Snca*Enh+37^-^) were created using a CRISPR Cas12a/AsCpf1 system, as we have done previously^1^. Single guide RNAs (sgRNAs; **TableS1**) were designed to flank the *Snca* enhancer interval and induce a 2.76 kb deletion between mm9 chr6: 60,742,326-60,745,103 (mm39 chr6: 60,769,316-60,772,093), 37.49 kb downstream from the *Snca* transcription start site (MGI: Tssr61200 – mm39 chr6:60,806,810-60,806,833). AsCpf1 scaffold sequences (5′-TAATTTCTACTCTTGTAGAT-3′) were ligated to the 5’ end of each sgRNA (Integrated DNA Technologies; IDT). Guides were diluted in Opti-MEM media (Thermo-Fisher Scientific # 31985070) to a final concentration of 2 uM, and CPF1 protein (IDT) was diluted in Opti-MEM media (Thermo-Fisher Scientific # 31985070) to a final concentration of 5 ng/μl. A total of 50 μl guide-CPF1 solution was electroporated into approximately 150 C57BL/6J × FVB/N F_1_ hybrid zygotes using a Nepa21 electroporator (Nepa Gene, Co. Ltd., Japan) with set pulse conditions as recommended by the manufacturer. Hybrid zygotes were allowed to rest for 30-60 minutes at 5% CO_2_, washed in M2 media (Sigma Aldrich # M7167), and implanted using standard embryo transfer surgery protocols^1^. The resulting founder mice were screened for deletion alleles using the flanking PCR primers (**Table S1; Figure S1**). Amplicons were Sanger sequenced to confirm the presence of the intended deletion. At sexual maturity, five founder animals and their offspring were backcrossed on a C57BL/6J (The Jackson Laboratory, Bar Harbor, Maine) background for 10 generations (N10) to generate congenic strains. One of the five founder lines (V9) was slower to reproduce, reaching N8 when the other lines (L1, L2, Y1, and Y7) reached N10, and was excluded from subsequent analysis in the interest of time.

### Mouse genotyping

Mouse genomic DNA was isolated from ≤ 20 mg of ear tissue, obtained from 1-month-old mice. Tissue was incubated in lysis buffer (50 uM KCl, 10 uM Tris HCl pH9.0, 0.1% Triton X-100) and 5 μl of proteinase K (18.7 mg/ml; Sigma Aldrich #3115879001) at 60°C for 3 hours, followed by a 95°C incubation for 10 minutes. Extracted DNA was purified using a DNA Clean & Concentrator™-5 kit (Zymo Research #D4067), eluted in 6 μl of elution buffer, and quantified using a Nanodrop. All genomic DNA samples were diluted to a concentration of 20 ng/μl using nuclease free water.

Mouse genotype was assessed by polymerase chain reaction (PCR) and agarose gel electrophoresis using custom primers designed to flank and bridge the enhancer locus (**Table S1; Figure S1**). A 25 μl reaction volume – consisting of 12.5 μl GoTaq® Green Master Mix (Promega #M7123); 1 μl each of forward, reverse, and internal primers at a concentration of 10 μM; 8.5 μl nuclease free water; and 1 μl of template DNA at a concentration of 20 ng/μl – was amplified using an initial denaturation step of 95°C for 5 minutes; followed by 30 cycles of 95°C for 30 seconds, 56°C for 30 seconds, and 72°C for 60 seconds; and a final extension step of 72°C for 5 minutes. Amplicon size was assessed on a 1% (w/v) agarose gel.

### Snca Quantification via Single Molecule Fluorescent in situ Hybridization

Single molecule fluorescent in situ hybridization (smFISH) was performed using the RNAscope® Multiplex Fluorescent Reagent Kit v2 Assay (ACDBio, user manual document 323100-USM/Rev Date: 02272019), following protocols for fixed-frozen tissue samples, with some modifications. All buffers, probes, and Opal™ dyes were prepared according to the ACD user manual.

First, N10 mice from each of the 4 founder lines (3 mice/genotype/line) were given 10 μg/g body weight of a 77.5% (v/v) saline, 15% (v/v) ketamine (100 mg/mL), 7.5% (v/v) xylazine (100 mg/mL) anesthesia cocktail via intraperitoneal (i.p.) injection. Next, they underwent transcardial perfusion with 4% (w/v) paraformaldehyde (PFA) solution composed of 40 g PFA (Sigma-Aldrich #P6148) in 100 uL of 10x PBS (pH 7.4) and 900 uL water treated with 0.1% (v/v) DEPC (Diethyl pyrocarbonate; VWR #E174-100G). Brains were dissected and post-fixed in 4% PFA solution overnight at 4°C. After 24 hours, brains were moved to a 30% (w/v) sucrose solution, where they remained for another 24 h or until the brain had sunk to the bottom of a 50 mL falcon tube. Brains were frozen using Cytocool™ II, Aerosol Freezing Spray (Epredia™ #8323), mounted on a microtome disk using O.C.T. Compound (Tissue-Tek® #4583), and placed in a microtome at −20°C. Midbrain coronal sections were cut to 12 μm thickness, and 10 sections at ∼120 μm intervals were transferred to Superfrost® Plus microscope slides (Fisherbrand™ #12-550-15). Slides were stored at −80°C until slide pre-treatment.

Prior to pre-treatment, slides were moved from −80°C to −20°C for 1 hour. Since brains were not mounted in O.C.T. Compound, the slides were next incubated for 1 hour in a HybEZ™ oven set to 37°C. Next, RNAscope® hydrogen peroxide treatment and target retrieval steps were performed as recommended by the manufacturer. Slides were dried at room temperature for 5 minutes before the hydrophobic barrier was applied and left to dry overnight at room temperature. The following day, RNAscope® Protease III reagent was applied to each slide, incubated for 30 minutes at 40°C, and then treated with the appropriate positive control probe, negative control probe, or probe mix (50 volumes of RNAscope™ Probe-Mm-Snca #313281: 1 volume RNAscope™ Probe-Mm-Th-C2 #317621-C2). All hybridization steps were performed as written. Meanwhile, Opal™ dyes (Akoya Biosciences; FP1487001KT and FP1488001KT), reconstituted in 75 μL of DMSO, were diluted according to our fluorophore optimization efforts. Opal™ 520 (Akoya Biosciences #FP1487001KT) was diluted 1:100 in TSA buffer (ACDBio) and Opal™ 570 (Akoya Biosciences #FP1488001KT) was diluted 1:1750 in TSA buffer (ACDBio). HRP-C1 and HRP-C2 signals were developed, as written, except that 300 μL of each Opal™ dye was added at the appropriate time. Finally, slides were counterstained using 250 μL of Hoechst 34580 (Thermo-Fisher Scientific #H21486) solution made from 3 μL Hoechst in 1.5 mL DEPC-PBS. Slides were incubated at room temperature for 10 minutes before the Hoechst stain was removed and 1-2 drops of ProLong Gold Antifade Mountant (Invitrogen™ #P36930) was added to each slide. A 24×50 mm glass coverslip (Fisherbrand™ #12-545-F) was carefully placed on each slide and slides were left to dry overnight in a cool, dark place.

Slides were imaged using a Nikon Eclipse Ti confocal microscope, with all imaging settings (offset, HV/gain, and laser percentage) normalized to Hoescht/DAPI signal. Images were exported in .tiff format at 16-bits/px. *Snca* expression was quantified using HALO software (indica labs; v3.3.2541.383), at a resolution of 0.41μm/px, in which DA neurons, defined as *Th-* and Hoechst/DAPI-expressing cells within the SN, were selected for inclusion in FISH scoring analysis. The area for analysis was manually defined, such that only DA neurons of the SN, and not adjacent brain regions (i.e., the ventral tegmental area), were included in analysis. *Snca* copy intensity was calculated for each slide image and was averaged over the right and left hemispheres for each animal (**Supplementary_Table_RNA_Quantification.xlsx;** Column AB). The geometric mean, standard deviation (SD), and standard error or the mean (SEM) of *Snca* copy intensity was calculated to include all three animals in each genotype group. Again, to determine if enhancer genotype significantly affects *Snca* expression, we employed one-way ANOVA and post-hoc Tukey HSD tests to calculate pair-wise significance for each founder line.

### Preparation and Stereotaxic Injection of α-synuclein Preformed Fibrils

Mouse recombinant full-length α-synuclein protein was purified as previously described, using the IPTG independent inducible pRK172 vector system^3^. α-syn PFF were also prepared as we have previously described^4–7^, by diluting endotoxin using the ToxinEraser™ Endotoxin Removal Kit (GenScript #L00338), then using a magnetic stirrer (1,000 r.p.m.) to mix 5 mg/mL α-Syn PFF in PBS at 37°C. The α-synuclein protein was incubated for a week before aggregates were diluted to 0.1 mg/mL in PBS and sonicated for 30 seconds (0.5 second pulse on/off) at 10% amplitude (Branson Digital sonifier, Danbury, CT, USA). α-Syn PFF was validated using atomic force microscopy and transmission electron microscopy and stored at −80 °C until stereotaxic injection.

At 3 months of age (postnatal day 90; P90), 10 mice/sex/genotype were randomly assigned to experimental (α-Syn PFF) or control (PBS) groups, and stereotaxic injections were performed as previously described^5–9^. All mice were anesthetized with 250 mg/kg of a 1.25% working solution of Tribromoethanol/Avertin made from 25g 2,2,2-Tribromoethanol powder (Sigma-Aldrich T48402) with 15.5ml 2-Methyl-2 butanol (Sigma-Aldrich 240486) via i.p. injection. A 26.5-gauge injection cannula was unilaterally inserted into the right hemisphere of the striatum guided by stereotaxic coordinates (mediolateral, 2.0 mm from bregma; anteroposterior, 0.2 mm; dorsoventral, 2.6 mm). Each mouse within the experimental and control groups was administered a 2 μL injection of α-Syn PFF (2.5 μg/μL in PBS) or PBS, respectively, at an infusion rate of 0.2 μL/minute. α-Syn PFF-induced PD pathology was allowed to develop for 6-months (180 days) following injection surgeries^5–10^, and by 9-months of age, all mice in the α-Syn PFF and PBS groups progressed to experimental endpoints consisting of motor, behavioural, and histopathological assays.

### Rotarod Test of Motor Control

All behaviour tests were performed in the Behavioral Core Facility at the Johns Hopkins University School of Medicine between 09:00–16:00 during the lights-on cycle. Rotarod tests were performed on 9-month-old mice, 6-months post α-Syn PFF/PBS injection in accordance with published methods^4,5,11^. Briefly, mice were acclimatized to the procedure room for 30 minutes before being placed on a rotamex V instrument equipped with photobeams and a sensor to automatically detect mice that fall from the rotarod. This rotarod cylinder slowly accelerated from 4 r.p.m. to 40 r.p.m. over 5 minutes. Rotamex settings remained constant throughout all trials: start speed, 4.0 r.p.m.; maximum speed, 40 r.p.m.; acceleration interval, 15 seconds; acceleration step, 2 r.p.m. The duration over which each animal remained on the rotarod was recorded (in seconds), and a trial ended when the animal fell off the rungs. All animals underwent three consecutive training days, each consisting of three trials. Next, over five test days, each mouse was given one warm-up run and one evaluated test run. The mean duration over which each mouse remained on the rotarod during their test run was calculated.

To determine if enhancer genotype, α-Syn PFF/PBS injection, or pairwise interactions of these variables significantly impacted motor function, we employed a two-way ANOVA, using R (v.4.3.3) function “aov”, and post-hoc Tukey HSD tests, using R function “TukeyHSD” specifying “conf.level=.95” to calculate pair-wise significance for each genotype and α-Syn PFF/PBS injection group. Outliers were removed such that data from n = 8-9 mice/sex/genotype/injection group (N = 106) were included in analysis.

### Open Field Test

Open field testing was performed on 9-month-old mice, 6-months post α-Syn PFF/PBS injection, as we have done previously^4,11^. In a quiet room, each mouse was placed against the wall of the open field arena – a rectangular plastic box (40 cm × 40 cm × 40 cm) divided into 36 (6 × 6) identical squares (6.6 cm × 6.6 cm), where the “central” sector was defined as the 4 central squares (2 × 2) and the “peripheral” sector was defined as the remaining squares. Each mouse was allowed to explore the arena for 15 minutes (900 seconds). Photobeam activity system (PAS) software connected to the open field equipment recorded the total number of beam breaks, which was used to determine gross locomotor activity of each mouse. Photobeam Activity System™ (PAS) software was used to measure ambulatory (A) and fine (F) movements at the center (cen) and periphery (per). Total center beam breaks were calculated by cenA+cenF; total peripheral beam breaks were calculated by perA+perF; locomotion was defined as total beam breaks (cenA+cenF+perA+perF); rears were defined as total number of vertical beam breaks; and anxiety was defined as the percentage of peripheral movements [perF+perA)/(cenA+cenF+perA+perF)*100].

To determine if enhancer genotype, α-Syn PFF/PBS injection, or pairwise interactions of these variables significantly impacted anxiety, locomotor, and exploratory behaviours, we employed two-way ANOVA and post-hoc Tukey HSD tests to calculate pair-wise significance for each genotype and α-Syn PFF/PBS injection group. Outliers were removed such that data from n = 9-10 mice/sex/genotype/injection group (N = 112) were included in analysis.

### Pole Descent Test of Motor Control

Pole descent tests were performed on 9-month-old mice, 6-months post α-Syn PFF/PBS injection as previously described^4,5^. Mice were placed on a gauze-wrapped metal rod 75 cm in height and 9 mm in diameter. Mice were placed on the top of the pole, facing down, and the total time (in seconds; 60 seconds maximum) taken for each mouse to initiate downward movement and reach the base of the pole with their front paws was recorded. All animals underwent three consecutive training days, each consisting of three trials, prior to four test days, during which mice were evaluated over two trials.

To determine if enhancer genotype, α-Syn PFF/PBS injection, or pairwise interactions of these variables significantly impacted motor control, we employed two-way ANOVA and post-hoc Tukey HSD tests to calculate pair-wise significance for each genotype and α-Syn PFF/PBS injection group. Outliers were removed such that data from n = 8-10 mice/sex/genotype/injection group (N = 114) were included in analysis.

### Grip Strength Test

Grip strength testing was performed on ≥9-month-old mice, ≥6-months post α-Syn PFF/PBS injection. Mice were acclimatized to the procedure room for 30 minutes before their forelimbs were placed on a specially designed grid connected to a Bioseb grip strength meter (BIO-GS4; v 3.47). Neuromuscular function was determined by the maximal peak force (g) required for an experimenter to pull the mouse off the grid by pulling each animal’s tail down at a constant speed. To reduce variability, one experimenter performed this assay for all mice. The grip strength meter was reset to 0.0 g before and after each test. All mice were naively evaluated over three trials.

To determine if enhancer genotype, α-Syn PFF/PBS injection, or pairwise interactions of these variables significantly impacted neuromuscular function and grip strength, we employed two-way ANOVA and post-hoc Tukey HSD tests to calculate pair-wise significance for each genotype and α-Syn PFF/PBS injection group. Outliers were removed such that data from n = 9-10 mice/sex/genotype/injection group (N = 118) were included in analysis.

### Immunohistochemistry and Immunofluorescence

Following the acquisition of all motor and behavioural data, 5 mice/sex/treatment/genotype were anesthetized with 250 mg/kg Tribromoethanol/Avertin by i.p. injection and underwent transcardial perfusion with 4% PFA (pH7.4; Boster #AR1068). Brains were dissected from each animal and post-fixed in 4% PFA overnight. The following day, brains were moved to a cryoprotectant 30% (w/v) sucrose solution, where they remained for 2-3 days or until the brain had sunk to the bottom of a 50 mL falcon tube. Next, brains were frozen using O.C.T. Compound, coronal sections were cut to 30 μm thickness using a microtome, and sections were placed, free-floating, in 9 wells of a 12-well tissue culture plate in a storage solution of 0.01 M PBS. Sections were stored at −20°C overnight prior to IHC staining.

Immunohistochemistry (IHC) was performed with 30µm thick brain sections. The sections were washed in PBS 3 times for 10 minutes before blocking in PBS containing 5% (v/v) goat serum (Jackson ImmunoResearch #005-000-121) and 0.3% (v/v) Triton™ X-100 (Millipore Sigma #T9284). After 1 h in blocking buffer, sections were washed in PBS 3 times for 10 minutes. Next, sections were incubated in 1:500 dilution of an anti-TH primary antibody (Novus Biologicals #NB300-109) and were left to shake gently overnight at 4°C. The next day, sections were washed in PBS 3 times for 10 minutes and then incubated with 1:250 dilution of biotin-conjugated anti-rabbit IgG secondary antibody (Vector laboratories #BA-1000) in PBS containing 0.3% Triton™ X-100 for 2h at RT. After washing with PBS again, brain sections were placed in ABC solution (Vector Laboratories #PK-6100) for 2 hours at room temperature (RT). After a final wash in PBS 3 times for 10 minutes, brain sections were developed using SigmaFast DAB peroxidase substrate (Millipore Sigma #D4293). The sections were mounted on a gelatin-coated slide, followed by counterstaining with Nissl (0.09% v/v thionin; Millipore Sigma, 861340-25G). Image analysis was performed using Stereo Investigator software (MicroBright-Field, VT, USA), including an Axiophot photomicroscope (Carl Zeiss) and Hitachi HV C20 camera. TH- and Nissl-positive dopaminergic neurons in SN region were counted by a blinded investigator.

For immunofluorescence, sections were incubated in 1:500 dilution of anti-pS129-α-syn antibodies (BioLegend #825701), anti-GFAP antibodies (Invitrogen #14-9892-82) and anti-Iba-1 antibodies (Wako #019-19741) overnight at 4°C. The next day, sections were incubated in 1:250 dilution of Alexa-Fluor 488- and 594-conjugated secondary antibodies (Invitrogen). The images were obtained by confocal microscopy (LSM 880, Carl Zeiss), using Zen software. Signal intensity and counting were conducted using ImageJ software (v1.48). GraphPad Prism (v10.3.0) was used to plot the data and perform a Two-way ANOVA (or Mixed-Model) with the following parameters: “Multiple comparisons test” = Tukey, “Multiple comparisons options” = Report multiplicity adjusted P value for each comparison, “Family-wise alpha threshold and confidence” = 0.05 (95% confidence interval).

### Tissue Lysate and Western Blot Analysis

After perfusion with cold 1x-PBS, nonionic detergent-soluble and -insoluble fractions were made by homogenization of tissue in soluble lysis buffer (50 mM Tris-HCl, pH 7.4; 150 mM NaCl;1% Triton x100; phosphatase inhibitor cocktail II and III (Sigma-Aldrich); and complete protease inhibitor; DW). The homogenate was centrifuged at 22,000 x g for 20 minutes at 4°C, and the resulting pellet (P1) and supernatant (S1, soluble part) fractions were collected. The P1 was washed 2 times in soluble lysis buffer, re-suspended using insoluble lysis buffer (50 mM Tris-HCl, pH 7.4; 150 mM NaCl; 1% Triton x100; 2% SDS; phosphatase inhibitor cocktail II and III (Sigma-Aldrich); and complete protease inhibitor; DW) and then sonicated (20% amplitude, 1 s pulse on/off, total 5s). The P1 was centrifuged at 22,000 x g for 20 minutes at room temperature, and the resulting pellet (P2) and supernatant (S2, insoluble part) fractions were collected. For western blot analysis, protein concentrations were measured using a Bicinchoninic Acid (BCA) assay (Pierce, Rockford, IL, USA). The protein samples were separated on 8 - 16% gradient SDS-PAGE gels and transferred to nitrocellulose (NC) membrane (0.45µm, Bio-Rad). After blocking with 5% non-fat milk or 3% BSA in TES-T (Tris-buffered saline with 0.1% tween-20) for 1h at room temperature, and incubated overnight at 4°C with anti-TH (1:1000, Novus Biologicals #NB300-109), anti-DAT (1:1000, Millipore Sigma #D6944), anti-pS129-a-syn (1:1000, Cell Signaling #23706), anti-a-syn (1:1000, Cell Signaling #2642S), anti-IBA1 (1:1000, Wako #019-19741), anti-GFAP (1:1000, Invitrogen #14-9892-82). Membranes were then washed and incubated with HRP-conjugated secondary antibodies (1:10000, Millipore #32230, #32260) for 1h at RT. Protein were detected using ECL solution and an Amersham Imager 800 (GE healthcare Life Sciences,). Results of western blot was quantified using ImageJ (v1.48). Plotting and statistical analysis was performed using GraphPad Prism (v10.3.0), as stated above.

---

**Figure S1.**
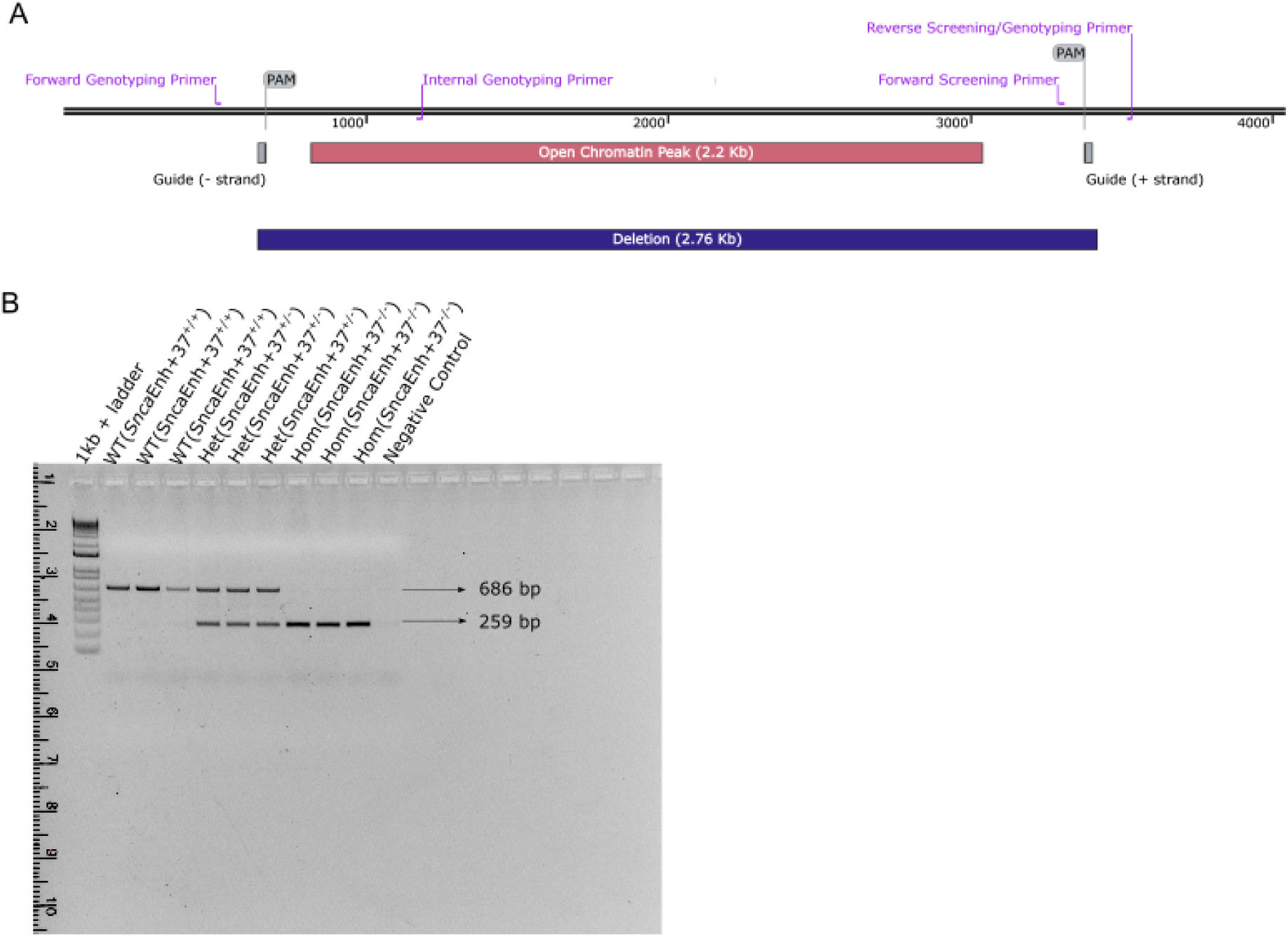
Characterization of *Snca*Enh+37 mice. **(A)** A SnapGene map showing the location of genotyping primers, screening primers, guide RNA sequences, *Snca*Enh+37 deletion, and their relative position to the lifted-over open chromatin peak identified in McClymont *et al* [*17*]. **(B)** A 1% agarose gel run with 3 replicates of each *Snca*Enh+37 mouse genotype: WT(*Snca*Enh+37^+/+^) = 686 bp amplicon between “Forward Genotyping Primer” and “Internal Genotyping Primer”; Hom(*Snca*Enh+37^-/-^) = 259 bp amplicon between “Forward Genotyping Primer” and “Reverse Screening/Genotyping Primer,” and Het(*Snca*Enh+37^+/-^) = 686 bp and 259 bp amplicon.

**Figure S2.**
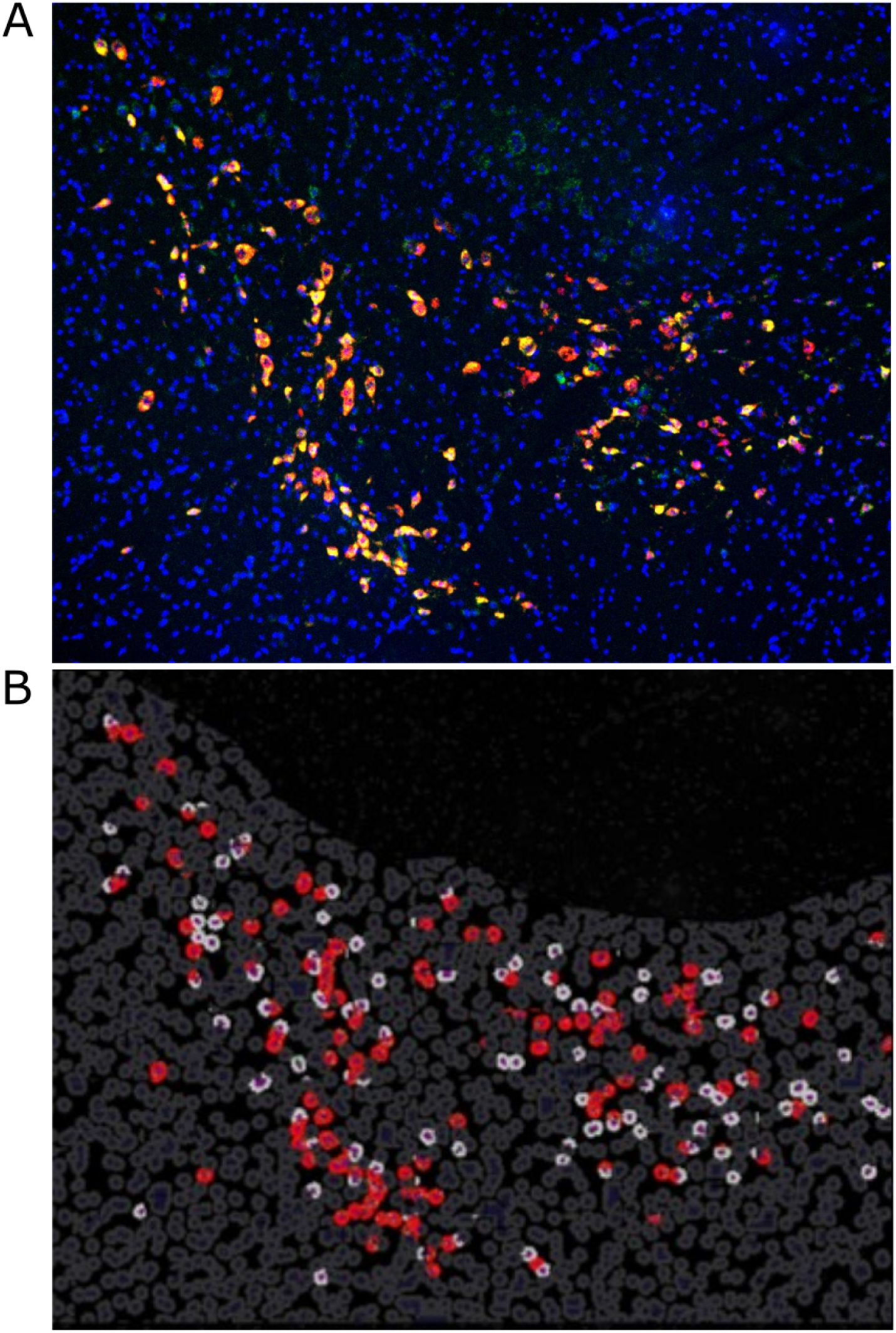
Representative images of (**A**) smFISH/RNAscope staining in the substantia nigra of a Het(*Snca*Enh+37^+/-^) founder mouse (L1); blue = Hoechst, red = *Th*, green = *Snca*, yellow = colocalization of *Th* and *Snca* and **(B)** the HALO logic gate used to identify dopaminergic neurons from RNAscope images (purple = nucleus, light gray = cytoplasm of *Th*+ cells, dark gray = cytoplasm of *Th*-cells, red = cytoplasm of Th+ cells containing *Snca* transcripts).

**Figure S3.**
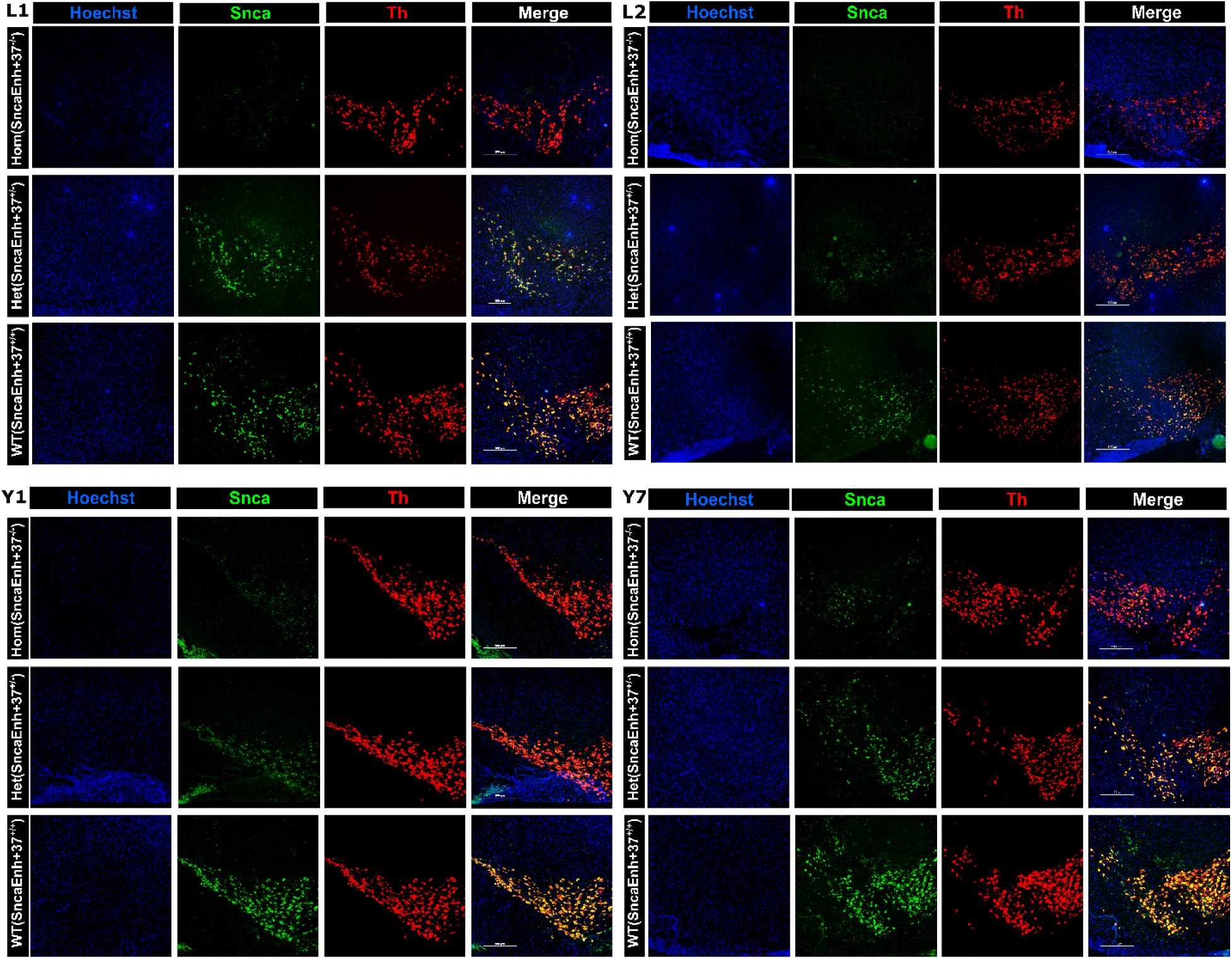
Representative smFISH/RNAscope images from *Snca*Enh+37 founder mice, showing WT(*Snca*Enh+37^+/+^), Het(*Snca*Enh+37^+/-^), and Hom*(Snca*Enh+37^-/-^) mice from founder lines L1, L2, Y1, and Y7. Scale bars = 100μm.

**Figure S4.**
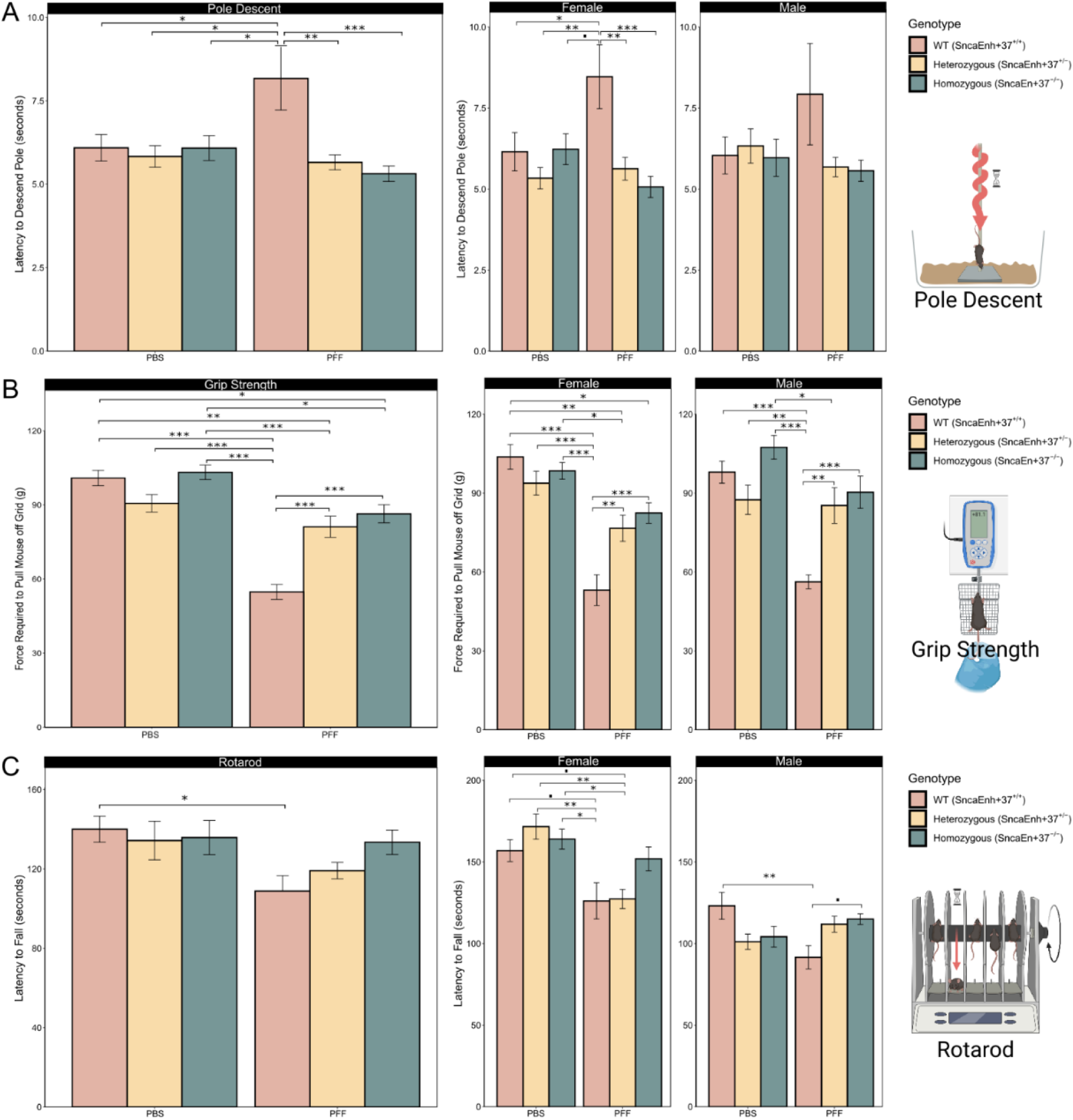
Mice lacking *Snca*Enh+37 are protected against motor deficits as measured by: **(A)** pole descent performance; **(B)** Grip strength; and **(C)** Rotarod performance. **(D)** Mice lacking *Snca*Enh+37 are also protected against anxiety-like behaviours. Bar charts report both combined and sex-dependent analyses of 8-10 mice/sex/genotype/treatment group. Error bars represent standard error (SE). Significance was determined using a two-way ANOVA and Tukey’s Honestly Significant Difference (HSD) test with a 95% confidence interval. See **Tables S3-S26** for ANOVA summary statistics. * = p < 0.05, ** = p < 0.01, *** = p < 0.001.

**Figure S5.**
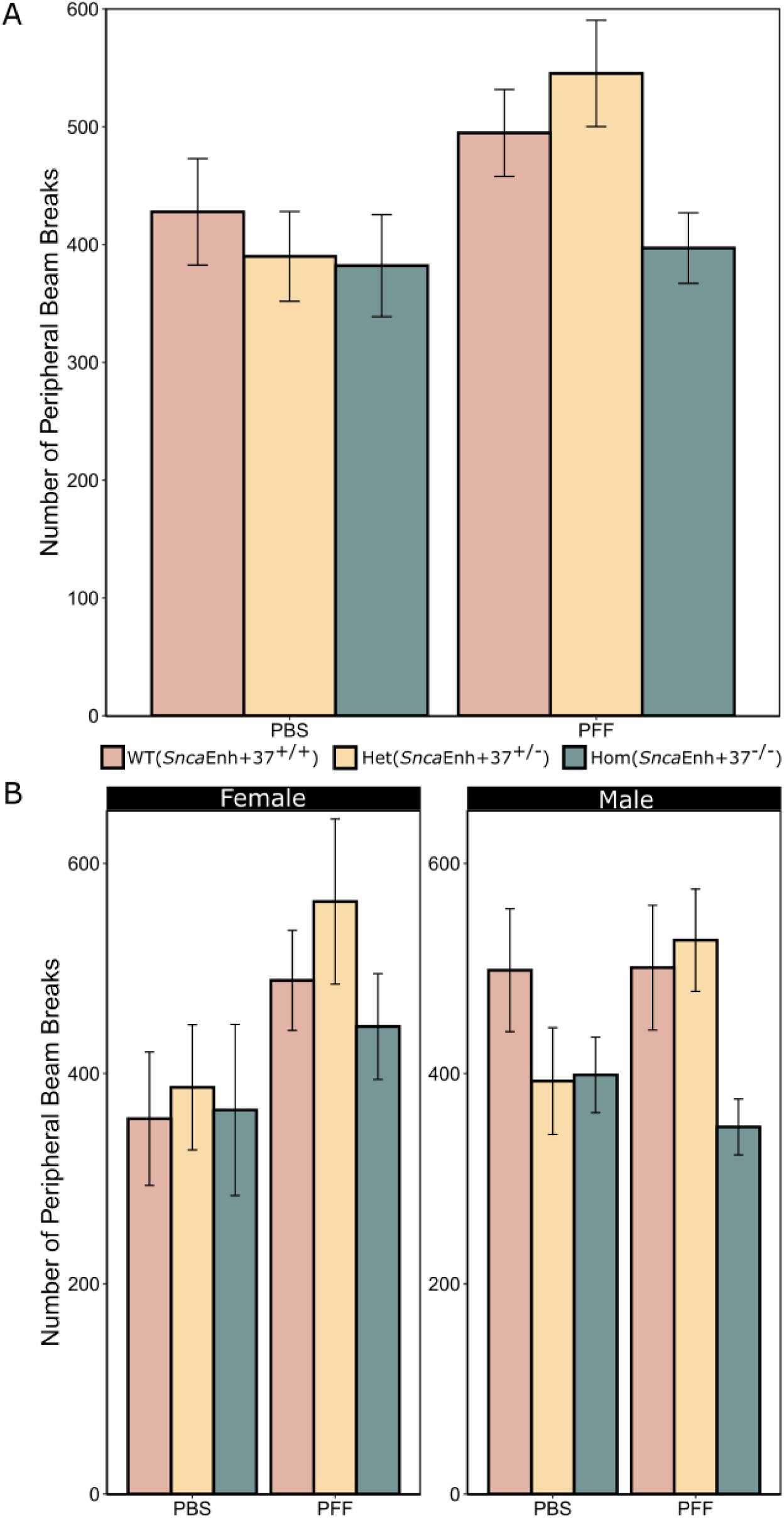
Mice lacking both copies of *Snca*Enh+37 record more peripheral movement. **(A)** Bar chart showing center movement for 16-20 mice/genotype/treatment group. **(B)** Bar chart showing sex differences in peripheral movement of 8-10 mice/sex/genotype/treatment group. Error bars represent standard error (SE). Significance was determined using a two-way ANOVA and Tukey’s Honestly Significant Difference (HSD) test with a 95% confidence interval. See **Tables S18-S20** for ANOVA summary statistics. ▪ = p < 0.1, * = p < 0.05, ** = p < 0.01, *** = p < 0.001.

**Figure S6.**
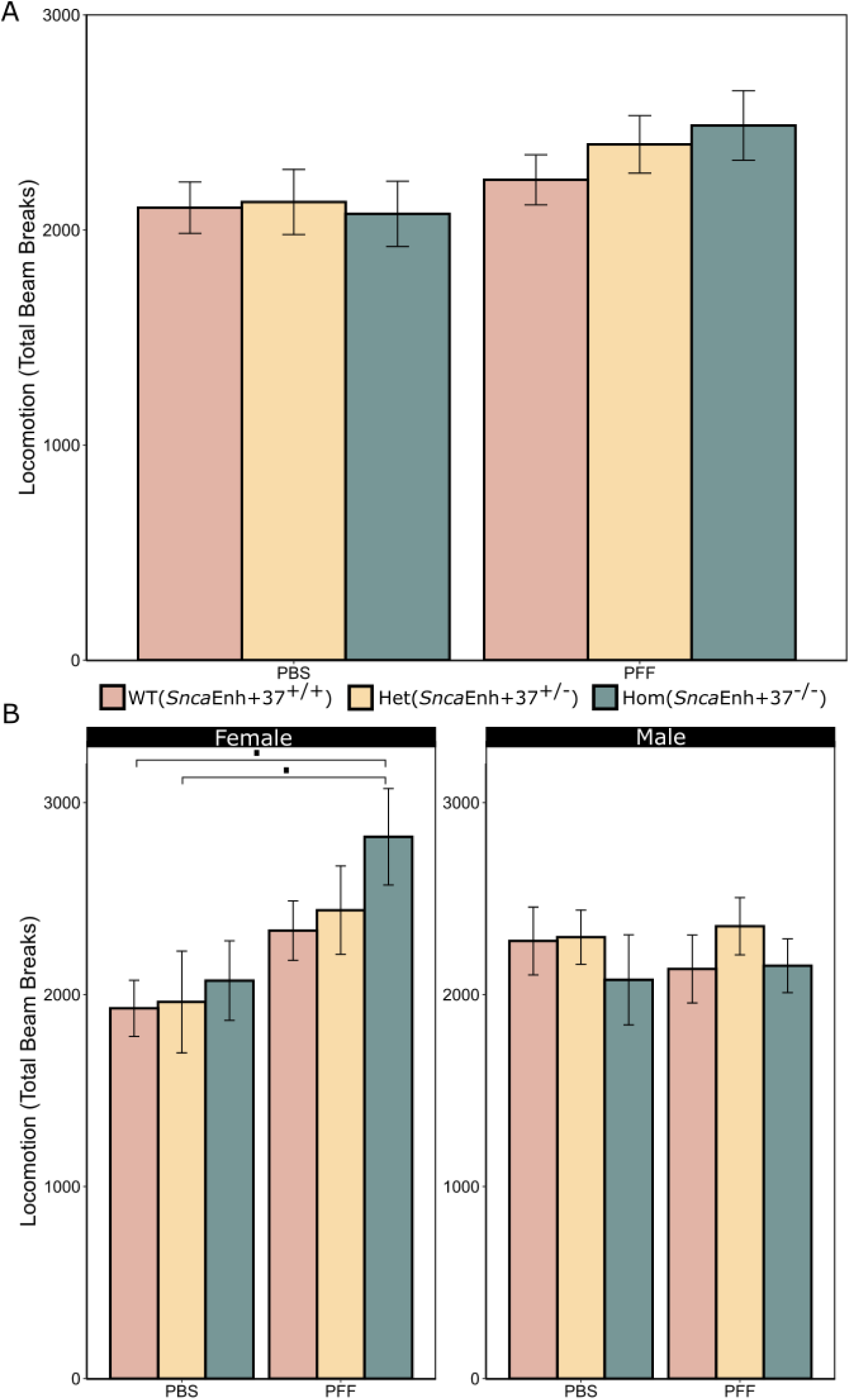
α-Syn PFF injected animals recorded more total movement in the open field chamber compared to PBS mice. **(A)** Bar chart showing locomotion for 16-20 mice/genotype/treatment group. **(B)** Bar chart showing sex differences in locomotion of 8-10 mice/sex/genotype/treatment group. Error bars represent standard error (SE). Significance was determined using a two-way ANOVA and Tukey’s Honestly Significant Difference (HSD) test with a 95% confidence interval. See **Tables S15-S17** for ANOVA summary statistics. ▪ = p < 0.1, * = p < 0.05, ** = p < 0.01, *** = p < 0.001.

**Figure S7.**
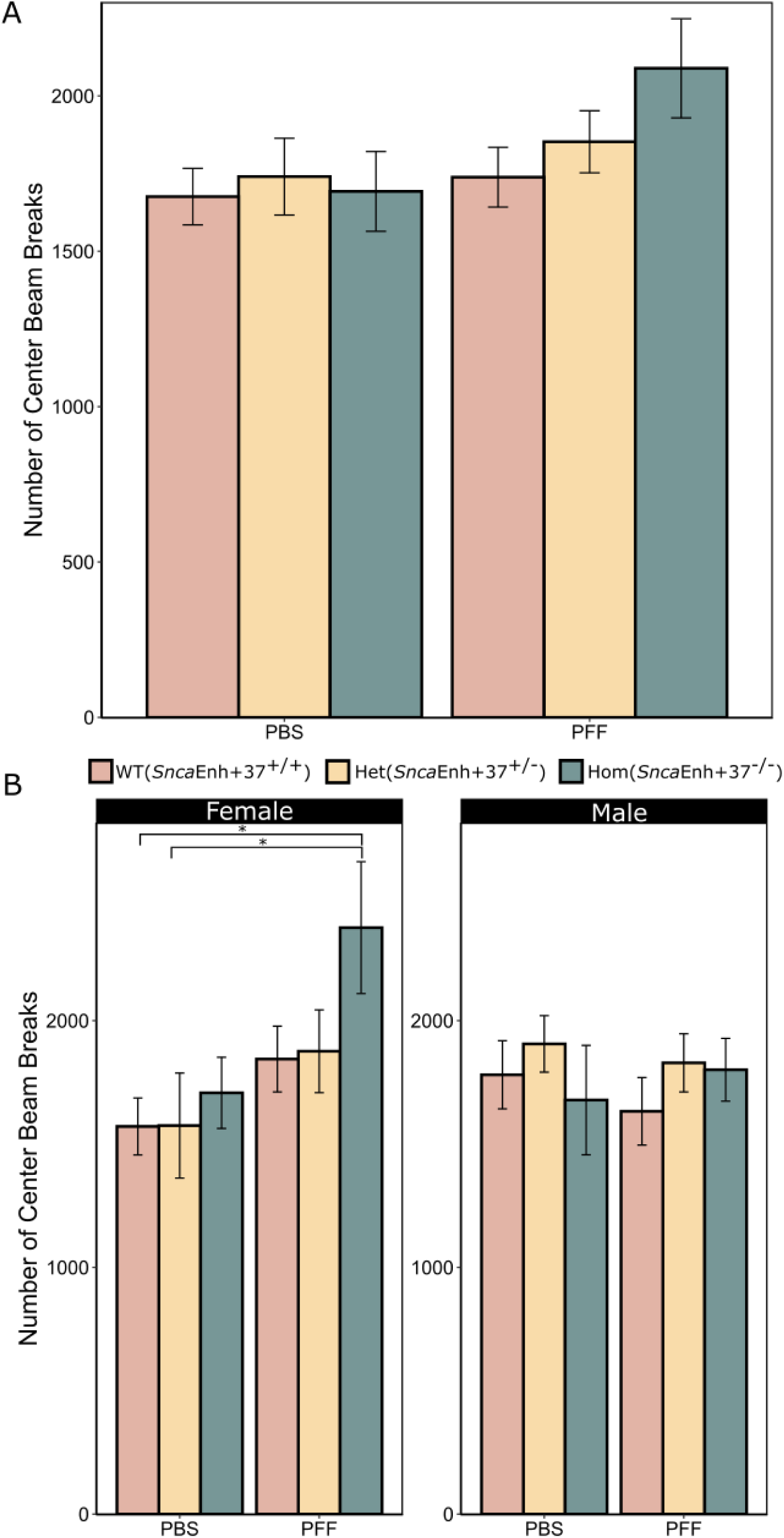
Mice lacking both copies of *Snca*Enh+37 record more center movement. **(A)** Bar chart showing center movement for 16-20 mice/genotype/treatment group. **(B)** Bar chart showing sex differences in center movement of 8-10 mice/sex/genotype/treatment group. Error bars represent standard error (SE). Significance was determined using a two-way ANOVA and Tukey’s Honestly Significant Difference (HSD) test with a 95% confidence interval. See **Tables S21-S23** for ANOVA summary statistics. ▪ = p < 0.1, * = p < 0.05, ** = p < 0.01, *** = p < 0.001.

**Figure S8.**
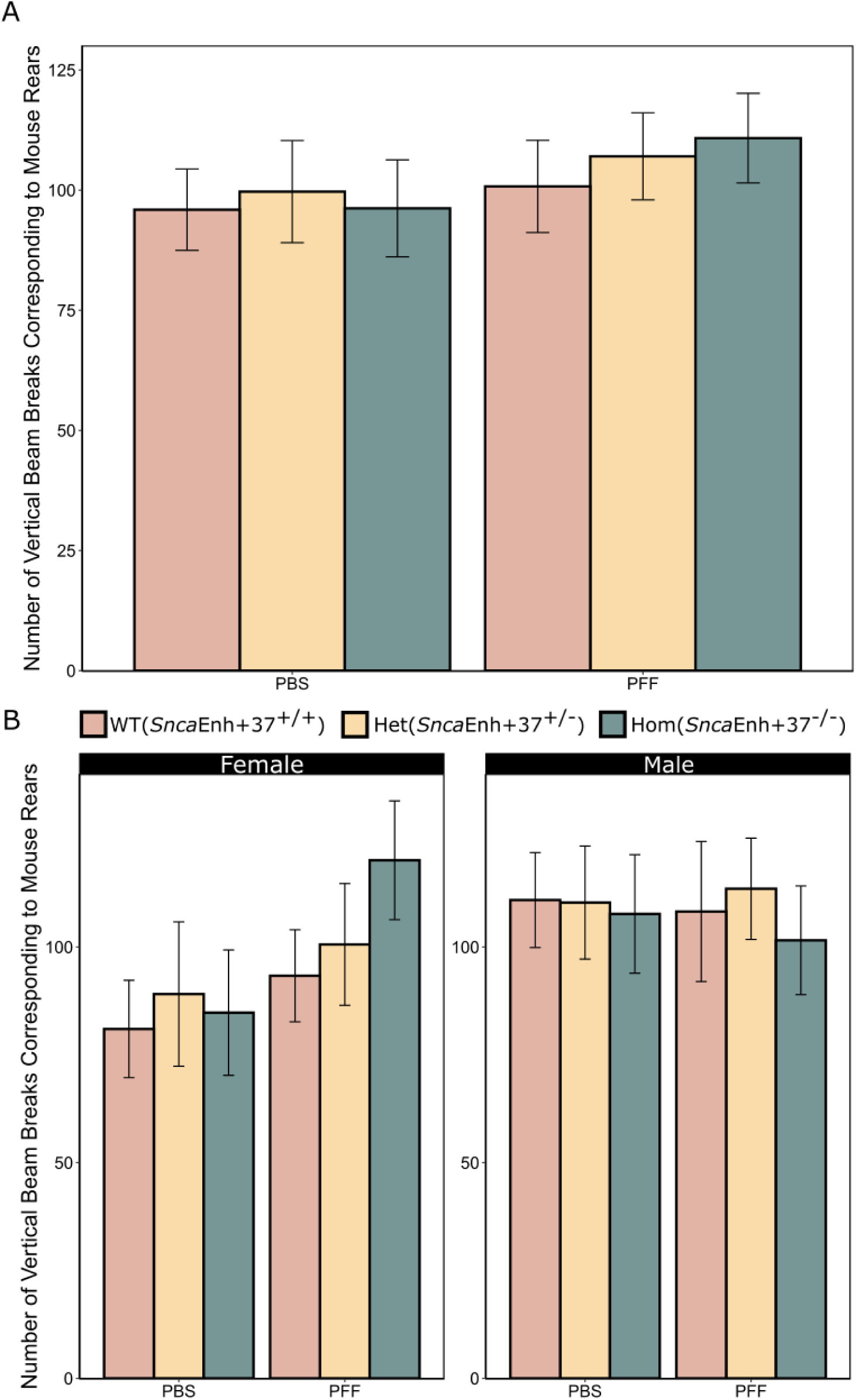
Mice lacking both copies of *Snca*Enh+37 show no differences in rearing behaviours. **(A)** Bar chart showing rearing behaviours for 16-20 mice/genotype/treatment group. **(B)** Bar chart showing sex differences in rearing behaviours of 8-10 mice/sex/genotype/treatment group. Error bars represent standard error (SE). Significance was determined using a two-way ANOVA and Tukey’s Honestly Significant Difference (HSD) test with a 95% confidence interval. See **Tables S24-S26** for ANOVA summary statistics. ▪ = p < 0.1, * = p < 0.05, ** = p < 0.01, *** = p < 0.001.

**Table S1.**
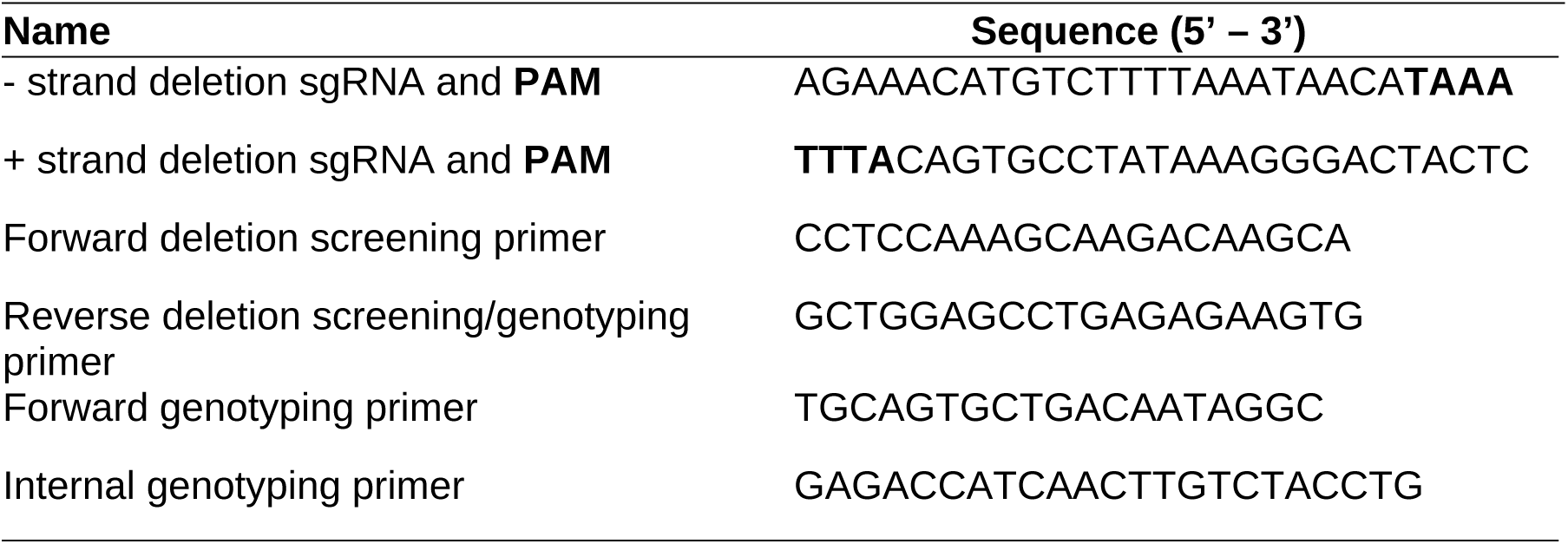
sgRNAs and primers used to generate and validate the *SncaEnh+37* mouse model.

**Table S2.** Summary statistics (ANOVA and Tukey’s HSD; CI = 95%) for *Snca* RNAscope.

| Founder Line | F-value | Pr(>F) | Pairwise Comparison | Difference (Tukey) | Adjusted p-value (Tukey) |
| --- | --- | --- | --- | --- | --- |
| L1 | 10.28 | 0.00251 | L1 WT-Het | 2.982879 | **0.0100273** |
|  |  |  | L1 Hom-Het | -0.536183 | 0.8007487 |
|  |  |  | L1 WT-Hom | 3.519062 | **0.0032181** |
| L2 | 15.22 | 0.000393 | L2 WT-Het | 1.699644 | **0.0007797** |
|  |  |  | L2 Hom-Het | 0.148493 | 0.9119488 |
|  |  |  | L2 WT-Hom | 1.551151 | **0.0016753** |
| Y1 | 5.852 | 0.0154 | Y1 WT-Het | 0.6431564 | 0.2551733 |
|  |  |  | Y1 Hom-Het | -0.6104493 | 0.1509333 |
|  |  |  | Y1 WT-Hom | 1.2536057 | **0.0118549** |
| Y7 | 5.313 | 0.0223 | Y7 WT-Het | 0.5982340 | 0.1541260 |
|  |  |  | Y7 Hom-Het | -0.3666896 | 0.4606378 |
|  |  |  | Y7 WT-Hom | 0.9649236 | **0.0183871** |

**Table S3.** Summary statistics (ANOVA and Tukey’s HSD; CI = 95%) for Pole Descent (All).

| **Pairwise Comparison** | **Difference (s)** | **Adjusted p-value (Tukey)** | **Significance** |
| --- | --- | --- | --- |
| Hom-Het | -0.04278 | 0.99588 | NS |
| **WT-Het** | **1.414557** | **0.011669** | **\*** |
| **WT-Hom** | **1.457337** | **0.011964** | **\*** |
| PFF-PBS | 0.397083 | 0.325389 | NS |
| Hom:PBS-Het:PBS | 0.250076 | 0.999234 | NS |
| WT:PBS-Het:PBS | 0.257602 | 0.999053 | NS |
| Het:PFF-Het:PBS | -0.17807 | 0.999824 | NS |
| Hom:PFF-Het:PBS | -0.51805 | 0.976903 | NS |
| **WT:PFF-Het:PBS** | **2.335813** | **0.011336** | **\*** |
| WT:PBS-Hom:PBS | 0.007526 | 1 | NS |
| Het:PFF-Hom:PBS | -0.42814 | 0.98967 | NS |
| Hom:PFF-Hom:PBS | -0.76813 | 0.894231 | NS |
| **WT:PFF-Hom:PBS** | **2.085736** | **0.041921** | **\*** |
| Het:PFF-WT:PBS | -0.43567 | 0.988058 | NS |
| Hom:PFF-WT:PBS | -0.77565 | 0.884589 | NS |
| **WT:PFF-WT:PBS** | **2.078211** | **0.038504** | **\*** |
| Hom:PFF-Het:PFF | -0.33998 | 0.996469 | NS |
| **WT:PFF-Het:PFF** | **2.513881** | **0.004207** | **\*\*** |
| **WT:PFF-Hom:PFF** | **2.853861** | **0.001274** | **\*\*** |

**Table s4.**
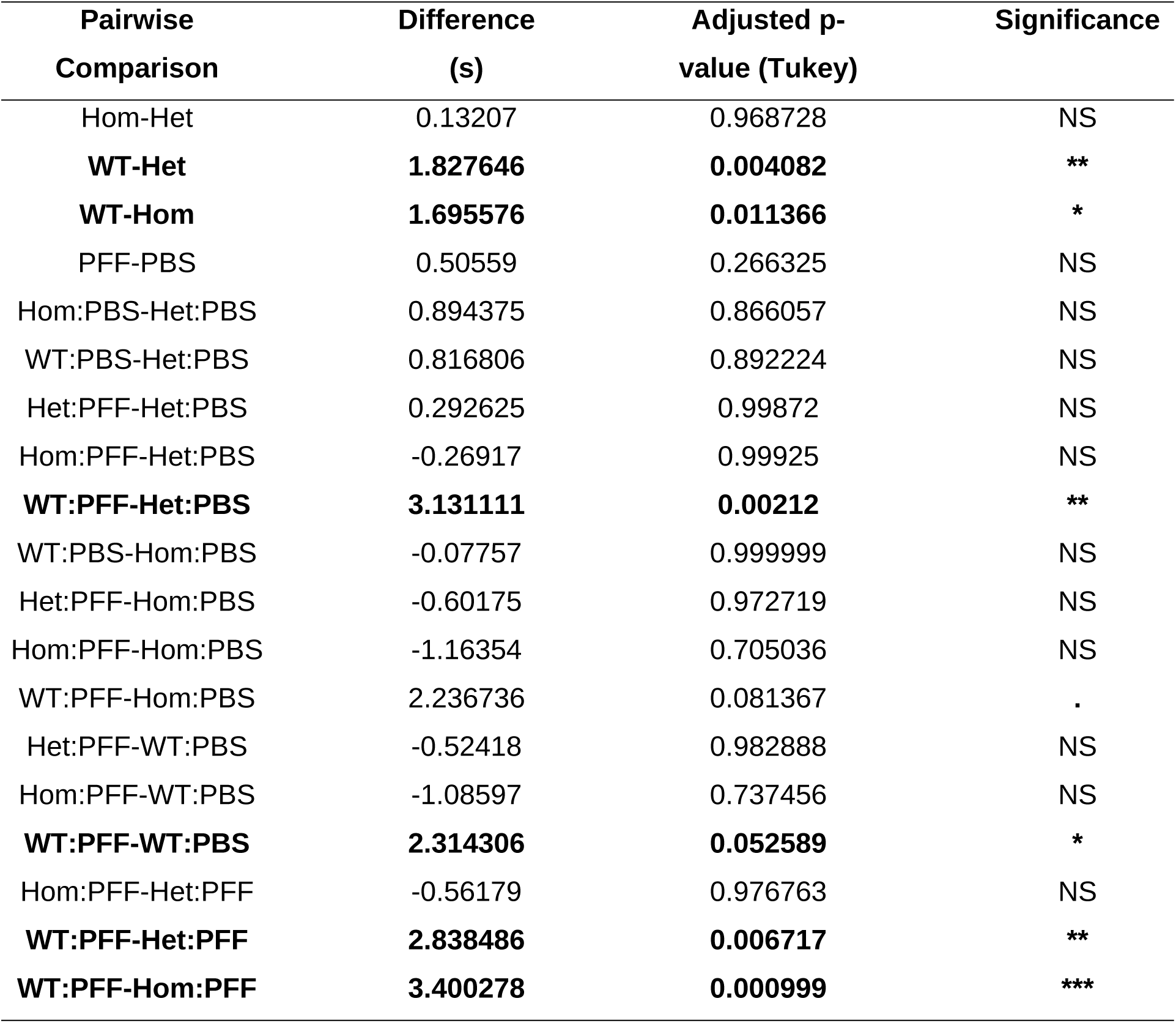
Summary statistics (ANOVA and Tukey’s HSD; CI = 95%) for Pole Descent (Female).

**Table S5.** Summary statistics (ANOVA and Tukey’s HSD; CI = 95%) for Pole Descent (Male).

| **Pairwise**<br>**Comparison** | **Difference (s)** | **Adjusted p-value**<br>**(Tukey)** | **Significance** |
| --- | --- | --- | --- |
| Hom-Het | -0.21425 | 0.963314 | NS |
| WT-Het | 1.035655 | 0.40579 | NS |
| WT-Hom | 1.249906 | 0.289731 | NS |
| PFF-PBS | 0.300732 | 0.653039 | NS |
| Hom:PBS-Het:PBS | -0.365 | 0.999568 | NS |
| WT:PBS-Het:PBS | -0.2955 | 0.999847 | NS |
| Het:PFF-Het:PBS | -0.65126 | 0.992352 | NS |
| Hom:PFF-Het:PBS | -0.76693 | 0.987174 | NS |
| WT:PFF-Het:PBS | 1.594534 | 0.723979 | NS |
| WT:PBS-Hom:PBS | 0.0695 | 1 | NS |
| Het:PFF-Hom:PBS | -0.28626 | 0.999853 | NS |
| Hom:PFF-Hom:PBS | -0.40193 | 0.999395 | NS |
| WT:PFF-Hom:PBS | 1.959534 | 0.520803 | NS |
| Het:PFF-WT:PBS | -0.35576 | 0.999573 | NS |
| Hom:PFF-WT:PBS | -0.47143 | 0.99869 | NS |
| WT:PFF-WT:PBS | 1.890034 | 0.56011 | NS |
| Hom:PFF-Het:PFF | -0.11567 | 0.999999 | NS |
| WT:PFF-Het:PFF | 2.245795 | 0.340564 | NS |
| WT:PFF-Hom:PFF | 2.361465 | 0.343267 | NS |

**Table S6.** Summary statistics (ANOVA and Tukey’s HSD; CI = 95%) for Grip Strength (All).

| **Pairwise Comparison** | **Difference (g)** | **Adjusted p-value (Tukey)** | **Significance** |
| --- | --- | --- | --- |
| **Hom-Het** | **8.791641** | **0.034177** | **\*** |
| WT-Het | -7.37674 | 0.090237 | . |
| **WT-Hom** | **-16.1684** | **3.42E-05** | **\*\*\*** |
| **PFF-PBS** | **-23.8648** | **1.84E-13** | **\*\*\*** |
| Hom:PBS-Het:PBS | 12.54921 | 0.12743 | NS |
| WT:PBS-Het:PBS | 10.24833 | 0.30326 | NS |
| Het:PFF-Het:PBS | -9.46516 | 0.378982 | NS |
| Hom:PFF-Het:PBS | -4.23167 | 0.955013 | NS |
| **WT:PFF-Het:PBS** | **-35.8806** | **1.06E-09** | **\*\*\*** |
| WT:PBS-Hom:PBS | -2.30088 | 0.997312 | NS |
| **Het:PFF-Hom:PBS** | **-22.0144** | **0.000265** | **\*\*\*** |
| **Hom:PFF-Hom:PBS** | **-16.7809** | **0.012947** | **\*** |
| **WT:PFF-Hom:PBS** | **-48.4298** | **5.06E-14** | **\*\*\*** |
| **Het:PFF-WT:PBS** | **-19.7135** | **0.001264** | **\*\*** |
| **Hom:PFF-WT:PBS** | **-14.48** | **0.044334** | **\*** |
| **WT:PFF-WT:PBS** | **-46.1289** | **7.22E-14** | **\*\*\*** |
| Hom:PFF-Het:PFF | 5.233492 | 0.889567 | NS |
| **WT:PFF-Het:PFF** | **-26.4155** | **6.43E-06** | **\*\*\*** |
| **WT:PFF-Hom:PFF** | **-31.6489** | **6.86E-08** | **\*\*\*** |

**Table S7.** Summary statistics (ANOVA and Tukey’s HSD; CI = 95%) for Grip Strength (Female).

| **Pairwise Comparison** | **Difference (g)** | **Adjusted p-value (Tukey)** | **Significance** |
| --- | --- | --- | --- |
| Hom-Het | 4.82886 | 0.545851 | NS |
| WT-Het | -5.44833 | 0.464025 | NS |
| WT-Hom | -10.2772 | 0.077566 | . |
| **PFF-PBS** | **-27.705** | **1.20E-09** | **\*\*\*** |
| Hom:PBS-Het:PBS | 4.680741 | 0.979504 | NS |
| WT:PBS-Het:PBS | 9.98 | 0.626359 | NS |
| Het:PFF-Het:PBS | -17.1567 | 0.095492 | . |
| Hom:PFF-Het:PBS | -11.3367 | 0.49053 | NS |
| **WT:PFF-Het:PBS** | **-40.7007** | **1.35E-06** | **\*\*\*** |
| WT:PBS-Hom:PBS | 5.299259 | 0.964892 | NS |
| **Het:PFF-Hom:PBS** | **-21.8374** | **0.019036** | **\*** |
| Hom:PFF-Hom:PBS | -16.0174 | 0.16152 | NS |
| **WT:PFF-Hom:PBS** | **-45.3815** | **1.89E-07** | **\*\*\*** |
| **Het:PFF-WT:PBS** | **-27.1367** | **0.0012** | **\*\*** |
| **Hom:PFF-WT:PBS** | **-21.3167** | **0.018537** | **\*** |
| **WT:PFF-WT:PBS** | **-50.6807** | **5.19E-09** | **\*\*\*** |
| Hom:PFF-Het:PFF | 5.82 | 0.941938 | NS |
| **WT:PFF-Het:PFF** | **-23.5441** | **0.009127** | **\*\*** |
| **WT:PFF-Hom:PFF** | **-29.3641** | **0.000574** | **\*\*\*** |

**Table S8.** Summary statistics (ANOVA and Tukey’s HSD; CI = 95%) for Grip Strength (Male).

| **Pairwise Comparison** | **Difference (g)** | **Adjusted p-value (Tukey)** | **Significance** |
| --- | --- | --- | --- |
| **Hom-Het** | **12.55492** | **0.049286** | **\*** |
| WT-Het | -9.21008 | 0.188646 | NS |
| **WT-Hom** | **-21.765** | **0.000354** | **\*\*\*** |
| **PFF-PBS** | **-20.0403** | **1.77E-05** | **\*\*\*** |
| Hom:PBS-Het:PBS | 19.94667 | 0.09571 | . |
| WT:PBS-Het:PBS | 10.51667 | 0.718939 | NS |
| Het:PFF-Het:PBS | -2.18576 | 0.999652 | NS |
| Hom:PFF-Het:PBS | 2.873333 | 0.998826 | NS |
| **WT:PFF-Het:PBS** | **-31.2267** | **0.001338** | **\*\*** |
| WT:PBS-Hom:PBS | -9.43 | 0.801243 | NS |
| **Het:PFF-Hom:PBS** | **-22.1324** | **0.039632** | **\*** |
| Hom:PFF-Hom:PBS | -17.0733 | 0.213938 | NS |
| **WT:PFF-Hom:PBS** | **-51.1733** | **8.87E-08** | **\*\*\*** |
| Het:PFF-WT:PBS | -12.7024 | 0.507564 | NS |
| Hom:PFF-WT:PBS | -7.64333 | 0.906794 | NS |
| **WT:PFF-WT:PBS** | **-41.7433** | **9.85E-06** | **\*\*\*** |
| Hom:PFF-Het:PFF | 5.059091 | 0.981668 | NS |
| **WT:PFF-Het:PFF** | **-29.0409** | **0.002553** | **\*\*** |
| **WT:PFF-Hom:PFF** | **-34.1** | **0.000371** | **\*\*\*** |

**Table S9.** Summary statistics (ANOVA and Tukey’s HSD; CI = 95%) for Rotarod (All).

| **Pairwise Comparison** | **Difference (s)** | **Adjusted p-value (Tukey)** | **Significance** |
| --- | --- | --- | --- |
| Hom-Het | 7.955829 | 0.533188 | NS |
| WT-Het | -1.36941 | 0.9811 | NS |
| WT-Hom | -9.32524 | 0.422679 | NS |
| **PFF-PBS** | **-16.3597** | **0.008033** | **\*\*** |
| Hom:PBS-Het:PBS | 1.523529 | 0.99999 | NS |
| WT:PBS-Het:PBS | 5.705948 | 0.99353 | NS |
| Het:PFF-Het:PBS | -15.1812 | 0.691176 | NS |
| Hom:PFF-Het:PBS | -0.86544 | 0.999999 | NS |
| WT:PFF-Het:PBS | -25.4592 | 0.163747 | NS |
| WT:PBS-Hom:PBS | 4.182418 | 0.998514 | NS |
| Het:PFF-Hom:PBS | -16.7047 | 0.597407 | NS |
| Hom:PFF-Hom:PBS | -2.38897 | 0.999917 | NS |
| WT:PFF-Hom:PBS | -26.9827 | 0.119549 | NS |
| Het:PFF-WT:PBS | -20.8871 | 0.330571 | NS |
| Hom:PFF-WT:PBS | -6.57139 | 0.988479 | NS |
| **WT:PFF-WT:PBS** | **-31.1651** | **0.040363** | **\*** |
| Hom:PFF-Het:PFF | 14.31574 | 0.753773 | NS |
| WT:PFF-Het:PFF | -10.278 | 0.925572 | NS |
| WT:PFF-Hom:PFF | -24.5938 | 0.207483 | NS |

**Table S10.**
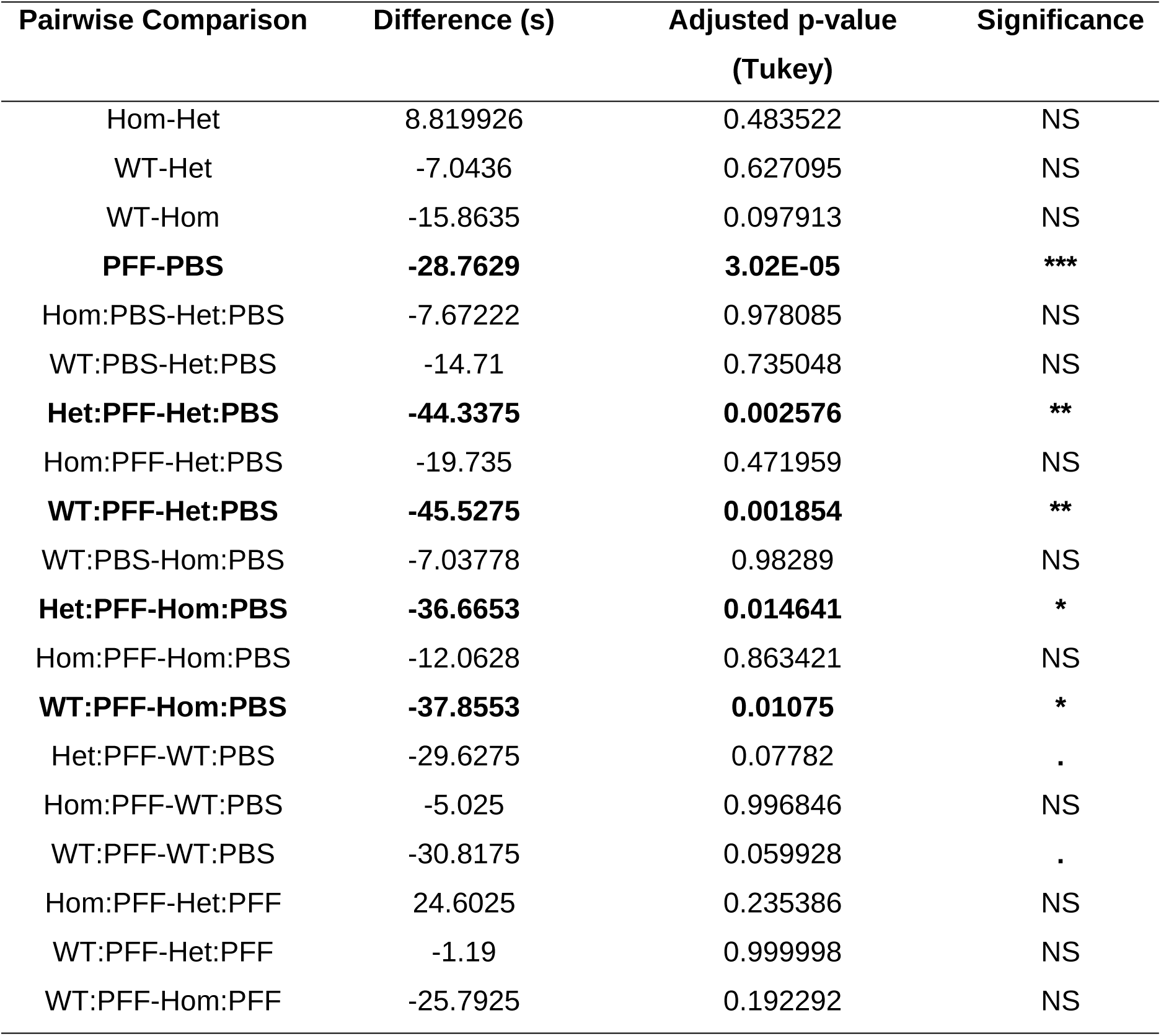
Summary statistics (ANOVA and Tukey’s HSD; CI = 95%) for Rotarod (Female).

**Table S11.** Summary statistics (ANOVA and Tukey’s HSD; CI = 95%) for Rotarod (Male).

| **Pairwise**<br>**Comparison** | **Difference**<br>**(s)** | **Adjusted p-**<br>**value (Tukey)** | **Significance** |
| --- | --- | --- | --- |
| Hom-Het | 3.086528 | 0.866191 | NS |
| WT-Het | 1.776013 | 0.951979 | NS |
| WT-Hom | -1.31051 | 0.975039 | NS |
| PFF-PBS | -3.29114 | 0.506983 | NS |
| Hom:PBS-Het:PBS | 3.053056 | 0.999178 | NS |
| WT:PBS-Het:PBS | 21.97333 | 0.10536 | NS |
| Het:PFF-Het:PBS | 10.73556 | 0.785356 | NS |
| Hom:PFF-Het:PBS | 13.85556 | 0.587929 | NS |
| WT:PFF-Het:PBS | -9.53944 | 0.871513 | NS |
| WT:PBS-Hom:PBS | 18.92028 | 0.250507 | NS |
| Het:PFF-Hom:PBS | 7.6825 | 0.944472 | NS |
| Hom:PFF-Hom:PBS | 10.8025 | 0.819891 | NS |
| WT:PFF-Hom:PBS | -12.5925 | 0.706633 | NS |
| Het:PFF-WT:PBS | -11.2378 | 0.75167 | NS |
| Hom:PFF-WT:PBS | -8.11778 | 0.930652 | NS |
| **WT:PFF-WT:PBS** | **-31.5128** | **0.007427** | **\*\*** |
| Hom:PFF-Het:PFF | 3.12 | 0.999087 | NS |
| WT:PFF-Het:PFF | -20.275 | 0.186779 | NS |
| WT:PFF-Hom:PFF | -23.395 | 0.103057 | . |

**Table S12.** Summary statistics (ANOVA and Tukey’s HSD; CI = 95%) for Anxiety (All).

| **Pairwise Comparison** | **Difference (% per)** | **Adjusted p-value (Tukey)** | **Significance** |
| --- | --- | --- | --- |
| Hom-Het | -2.77995 | 0.07035 | . |
| WT-Het | 0.732143 | 0.826848 | NS |
| **WT-Hom** | **3.512094** | **0.019094** | **\*** |
| PFF-PBS | 1.556611 | 0.131559 | NS |
| Hom:PBS-Het:PBS | 0.179018 | 0.999998 | NS |
| WT:PBS-Het:PBS | 1.640543 | 0.937448 | NS |
| Het:PFF-Het:PBS | 4.042777 | 0.180503 | NS |
| Hom:PFF-Het:PBS | -1.69614 | 0.92845 | NS |
| WT:PFF-Het:PBS | 3.866521 | 0.248472 | NS |
| WT:PBS-Hom:PBS | 1.461524 | 0.96531 | NS |
| Het:PFF-Hom:PBS | 3.863759 | 0.249198 | NS |
| Hom:PFF-Hom:PBS | -1.87516 | 0.904029 | NS |
| WT:PFF-Hom:PBS | 3.687502 | 0.326512 | NS |
| Het:PFF-WT:PBS | 2.402234 | 0.748124 | NS |
| Hom:PFF-WT:PBS | -3.33669 | 0.440611 | NS |
| WT:PFF-WT:PBS | 2.225978 | 0.820145 | NS |
| **Hom:PFF-Het:PFF** | **-5.73892** | **0.018305** | **\*** |
| WT:PFF-Het:PFF | -0.17626 | 0.999999 | NS |
| **WT:PFF-Hom:PFF** | **5.562663** | **0.030865** | **\*** |

**Table S13.** Summary statistics (ANOVA and Tukey’s HSD; CI = 95%) for Anxiety (All).

| **Pairwise Comparison** | **Difference (% per.)** | **Adjusted p-value (Tukey)** | **Significance** |
| --- | --- | --- | --- |
| **Hom-Het** | **-4.42821** | **0.050924** | **\*** |
| WT-Het | -1.60125 | 0.661188 | NS |
| WT-Hom | 2.826962 | 0.300528 | NS |
| PFF-PBS | 1.843285 | 0.228847 | NS |
| Hom:PBS-Het:PBS | -3.07552 | 0.843111 | NS |
| WT:PBS-Het:PBS | -1.7003 | 0.986078 | NS |
| Het:PFF-Het:PBS | 2.649198 | 0.899716 | NS |
| Hom:PFF-Het:PBS | -3.1317 | 0.832873 | NS |
| WT:PFF-Het:PBS | 1.146995 | 0.997768 | NS |
| WT:PBS-Hom:PBS | 1.375227 | 0.995337 | NS |
| Het:PFF-Hom:PBS | 5.724721 | 0.255677 | NS |
| Hom:PFF-Hom:PBS | -0.05618 | 1 | NS |
| WT:PFF-Hom:PBS | 4.222518 | 0.613805 | NS |
| Het:PFF-WT:PBS | 4.349494 | 0.556128 | NS |
| Hom:PFF-WT:PBS | -1.43141 | 0.994377 | NS |
| WT:PFF-WT:PBS | 2.847291 | 0.892072 | NS |
| Hom:PFF-Het:PFF | -5.7809 | 0.246019 | NS |
| WT:PFF-Het:PFF | -1.5022 | 0.992081 | NS |
| WT:PFF-Hom:PFF | 4.278698 | 0.60033 | NS |

**Table S14.** Summary statistics (ANOVA and Tukey’s HSD; CI = 95%) for Anxiety (Male).

| **Pairwise**<br>**Comparison** | **Difference (Tukey)** | **Adjusted p-value**<br>**(Tukey)** | **Significance** |
| --- | --- | --- | --- |
| Hom-Het | -1.13169 | 0.778292 | NS |
| WT-Het | 3.065536 | 0.169705 | NS |
| **WT-Hom** | **4.197225** | **0.046701** | **\*** |
| PFF-PBS | 1.269938 | 0.360665 | NS |
| Hom:PBS-Het:PBS | 3.43356 | 0.696183 | NS |
| WT:PBS-Het:PBS | 4.981381 | 0.301467 | NS |
| Het:PFF-Het:PBS | 5.436356 | 0.190231 | NS |
| Hom:PFF-Het:PBS | -0.26058 | 0.999998 | NS |
| WT:PFF-Het:PBS | 6.586046 | 0.077018 | . |
| WT:PBS-Hom:PBS | 1.547821 | 0.987565 | NS |
| Het:PFF-Hom:PBS | 2.002796 | 0.957124 | NS |
| Hom:PFF-Hom:PBS | -3.69414 | 0.652348 | NS |
| WT:PFF-Hom:PBS | 3.152486 | 0.784487 | NS |
| Het:PFF-WT:PBS | 0.454974 | 0.999961 | NS |
| Hom:PFF-WT:PBS | -5.24196 | 0.275067 | NS |
| WT:PFF-WT:PBS | 1.604665 | 0.98537 | NS |
| Hom:PFF-Het:PFF | -5.69694 | 0.173426 | NS |
| WT:PFF-Het:PFF | 1.149691 | 0.996465 | NS |
| WT:PFF-Hom:PFF | 6.846629 | 0.070648 | . |

**Table S15.** Summary statistics (ANOVA and Tukey’s HSD; CI = 95%) for Peripheral Movement (All).

| Pairwise Comparison | Difference (Tukey) | Adjusted p-value (Tukey) | Significance |
| --- | --- | --- | --- |
| Hom-Het | -78.0722 | 0.129798 | NS |
| WT-Het | -6.37778 | 0.986104 | NS |
| WT-Hom | 71.69444 | 0.193167 | NS |
| **PFF-PBS** | **81.83929** | **0.014503** | **\*** |
| Hom:PBS-Het:PBS | -7.84444 | 0.999993 | NS |
| WT:PBS-Het:PBS | 37.82222 | 0.985037 | NS |
| Het:PFF-Het:PBS | 155.4 | 0.061933 | . |
| Hom:PFF-Het:PBS | 7.1 | 0.999996 | NS |
| WT:PFF-Het:PBS | 104.8222 | 0.437607 | NS |
| WT:PBS-Hom:PBS | 45.66667 | 0.969303 | NS |
| Het:PFF-Hom:PBS | 163.2444 | 0.052539 | . |
| Hom:PFF-Hom:PBS | 14.94444 | 0.999843 | NS |
| WT:PFF-Hom:PBS | 112.6667 | 0.384033 | NS |
| Het:PFF-WT:PBS | 117.5778 | 0.307242 | NS |
| Hom:PFF-WT:PBS | -30.7222 | 0.994887 | NS |
| WT:PFF-WT:PBS | 67 | 0.857495 | NS |
| Hom:PFF-Het:PFF | -148.3 | 0.101459 | NS |
| WT:PFF-Het:PFF | -50.5778 | 0.947305 | NS |
| WT:PFF-Hom:PFF | 97.72222 | 0.546275 | NS |

**Table S16.**
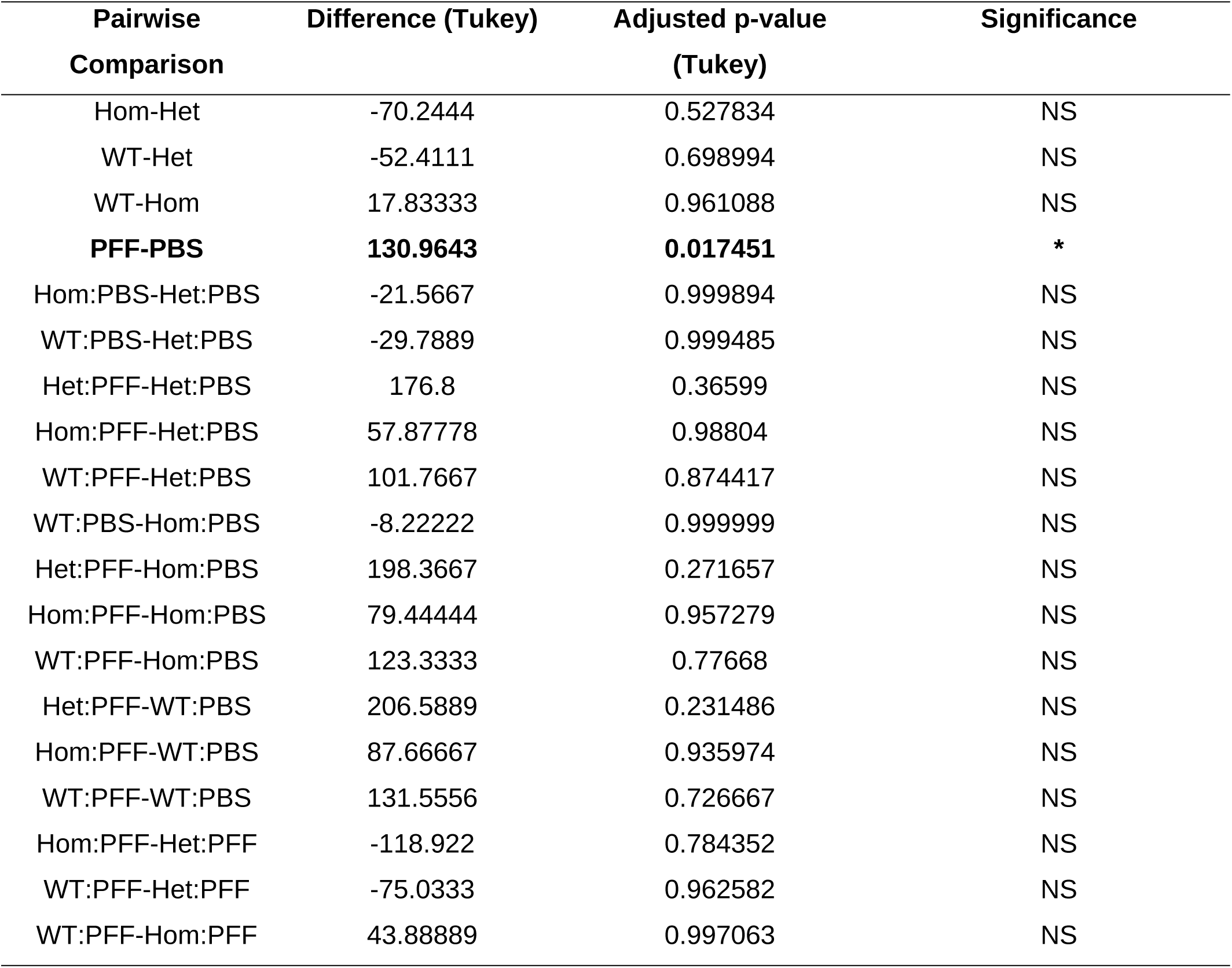
Summary statistics (ANOVA and Tukey’s HSD; CI = 95%) for Peripheral Movement (Female).

**Table S17.** Summary statistics (ANOVA and Tukey’s HSD; CI = 95%) for Peripheral Movement (Female).

| Pairwise Comparison | Difference (Tukey) | Adjusted p-value (Tukey) | Significance |
| --- | --- | --- | --- |
| Hom-Het | -85.9 | 0.182663 | NS |
| WT-Het | 39.65556 | 0.688011 | NS |
| **WT-Hom** | **125.5556** | **0.036114** | **\*** |
| PFF-PBS | 32.71429 | 0.410691 | NS |
| Hom:PBS-Het:PBS | 5.877778 | 0.999999 | NS |
| WT:PBS-Het:PBS | 105.4333 | 0.630967 | NS |
| Het:PFF-Het:PBS | 134 | 0.340057 | NS |
| Hom:PFF-Het:PBS | -43.6778 | 0.986954 | NS |
| WT:PFF-Het:PBS | 107.8778 | 0.607957 | NS |
| WT:PBS-Hom:PBS | 99.55556 | 0.708041 | NS |
| Het:PFF-Hom:PBS | 128.1222 | 0.42015 | NS |
| Hom:PFF-Hom:PBS | -49.5556 | 0.979558 | NS |
| WT:PFF-Hom:PBS | 102 | 0.686602 | NS |
| Het:PFF-WT:PBS | 28.56667 | 0.998203 | NS |
| Hom:PFF-WT:PBS | -149.111 | 0.2823 | NS |
| WT:PFF-WT:PBS | 2.444444 | 1 | NS |
| Hom:PFF-Het:PFF | -177.678 | 0.111042 | NS |
| WT:PFF-Het:PFF | -26.1222 | 0.99883 | NS |
| WT:PFF-Hom:PFF | 151.5556 | 0.26566 | NS |

**Table S18.** Summary statistics (ANOVA and Tukey’s HSD; CI = 95%) for Locomotion (All).

| **Pairwise**<br>**Comparison** | **Difference (Tukey)** | **Adjusted p-value**<br>**(Tukey)** | **Significance** |
| --- | --- | --- | --- |
| Hom-Het | 16.35 | 0.992439 | NS |
| WT-Het | -95.5667 | 0.772266 | NS |
| WT-Hom | -111.917 | 0.714375 | NS |
| **PFF-PBS** | **269.375** | **0.020625** | **\*** |
| Hom:PBS-Het:PBS | -55.4333 | 0.999756 | NS |
| WT:PBS-Het:PBS | -26.5444 | 0.999994 | NS |
| Het:PFF-Het:PBS | 267.6 | 0.729864 | NS |
| Hom:PFF-Het:PBS | 355.7333 | 0.466919 | NS |
| WT:PFF-Het:PBS | 103.0111 | 0.995164 | NS |
| WT:PBS-Hom:PBS | 28.88889 | 0.999991 | NS |
| Het:PFF-Hom:PBS | 323.0333 | 0.574703 | NS |
| Hom:PFF-Hom:PBS | 411.1667 | 0.330586 | NS |
| WT:PFF-Hom:PBS | 158.4444 | 0.969743 | NS |
| Het:PFF-WT:PBS | 294.1444 | 0.669705 | NS |
| Hom:PFF-WT:PBS | 382.2778 | 0.413551 | NS |
| WT:PFF-WT:PBS | 129.5556 | 0.987613 | NS |
| Hom:PFF-Het:PFF | 88.13333 | 0.997688 | NS |
| WT:PFF-Het:PFF | -164.589 | 0.960262 | NS |
| WT:PFF-Hom:PFF | -252.722 | 0.811046 | NS |

**Table S19.**
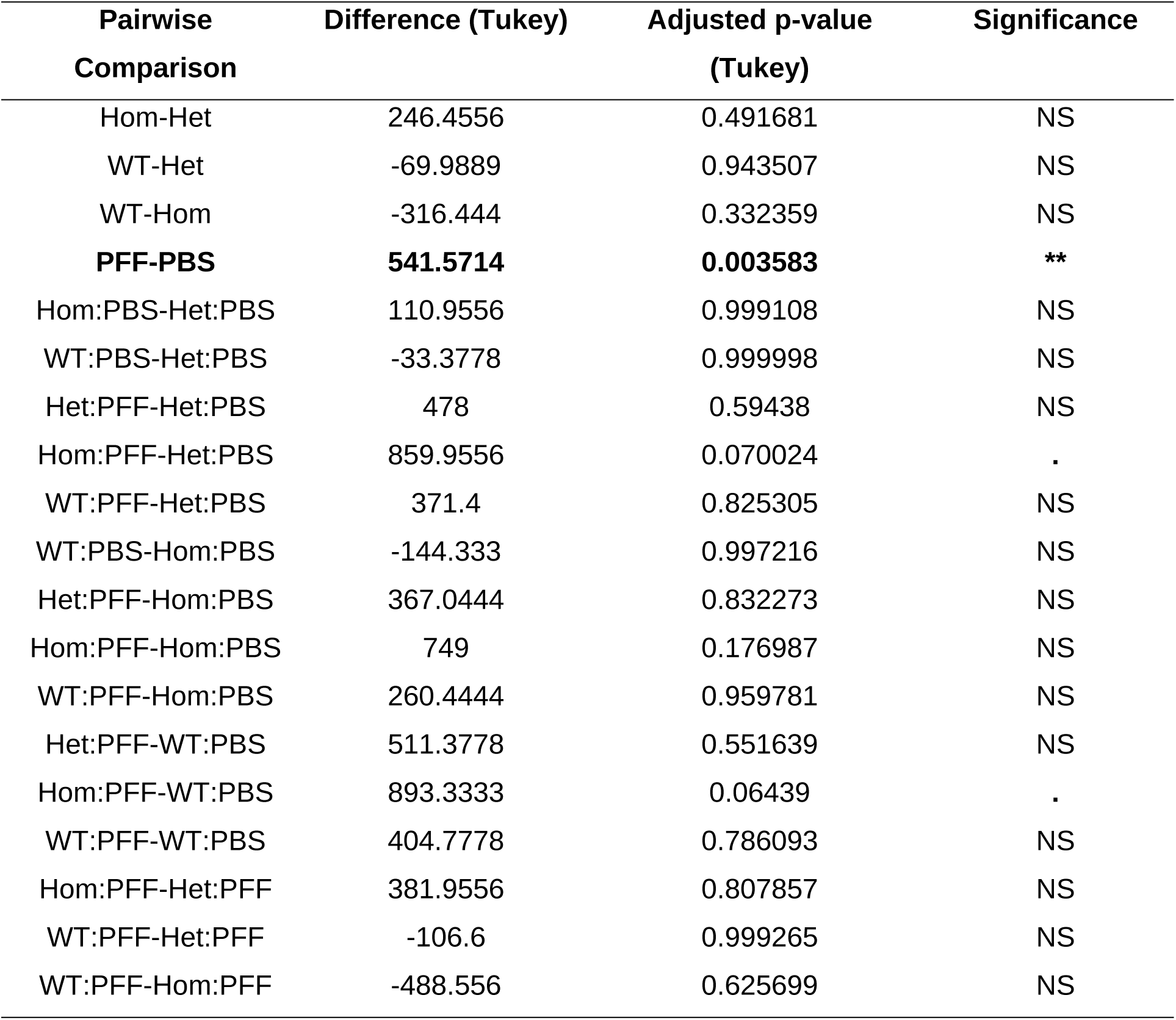
Summary statistics (ANOVA and Tukey’s HSD; CI = 95%) for Locomotion.

**Table S20.**
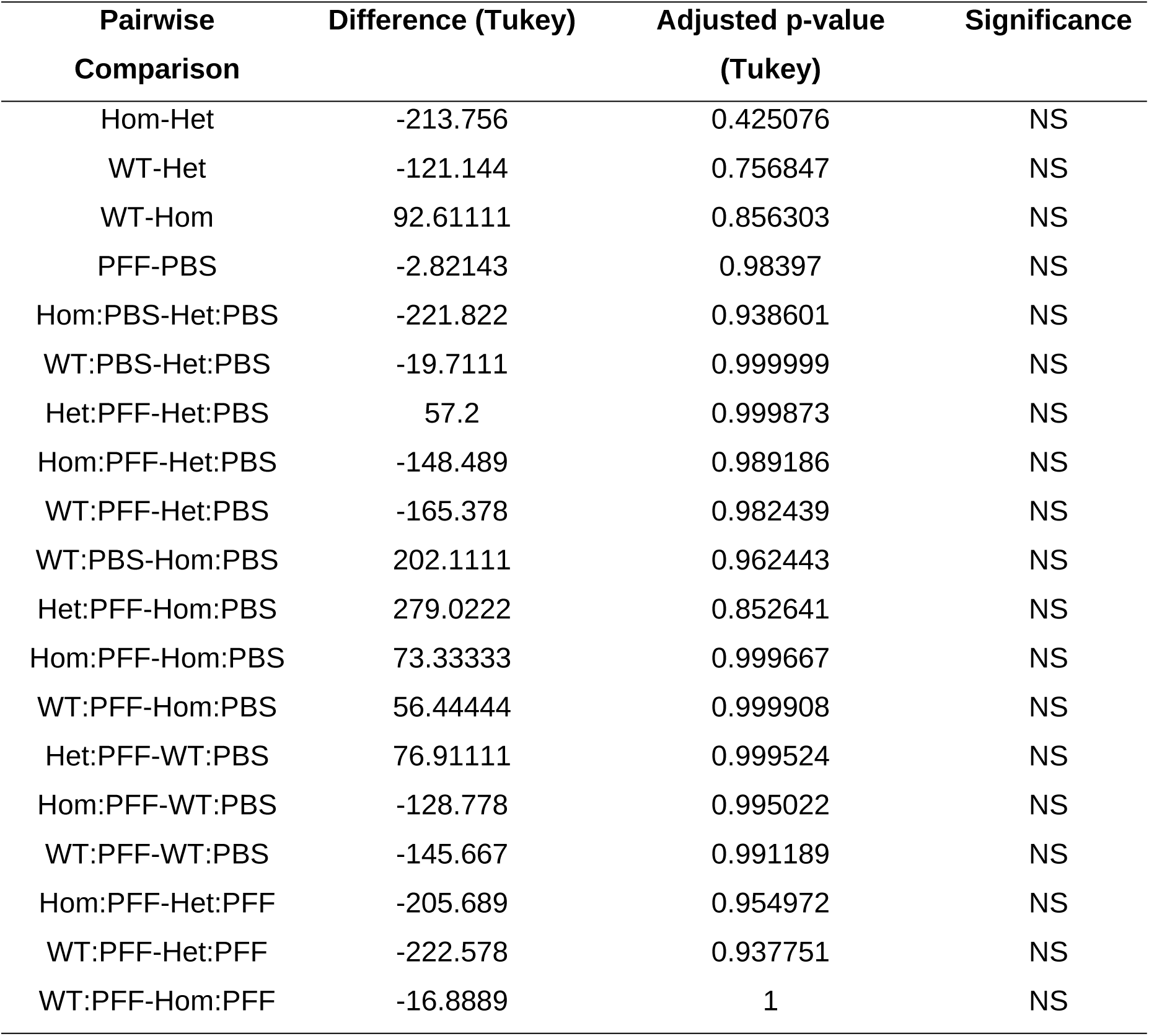
Summary statistics (ANOVA and Tukey’s HSD; CI = 95%) for Locomotion (Male).

**Table S21.** Summary statistics (ANOVA and Tukey’s HSD; CI = 95%) for Center Movement (All).

| **Pairwise**<br>**Comparison** | **Difference (Tukey)** | **Adjusted p-value**<br>**(Tukey)** | **Significance** |
| --- | --- | --- | --- |
| Hom-Het | 94.42222 | 0.702231 | NS |
| WT-Het | -89.1889 | 0.729385 | NS |
| WT-Hom | -183.611 | 0.285121 | NS |
| PFF-PBS | 187.5357 | 0.055302 | . |
| Hom:PBS-Het:PBS | -47.5889 | 0.999735 | NS |
| WT:PBS-Het:PBS | -64.3667 | 0.998847 | NS |
| Het:PFF-Het:PBS | 112.2 | 0.982389 | NS |
| Hom:PFF-Het:PBS | 348.6333 | 0.29761 | NS |
| WT:PFF-Het:PBS | -1.81111 | 1 | NS |
| WT:PBS-Hom:PBS | -16.7778 | 0.999999 | NS |
| Het:PFF-Hom:PBS | 159.7889 | 0.929278 | NS |
| Hom:PFF-Hom:PBS | 396.2222 | 0.194835 | NS |
| WT:PFF-Hom:PBS | 45.77778 | 0.999807 | NS |
| Het:PFF-WT:PBS | 176.5667 | 0.895485 | NS |
| Hom:PFF-WT:PBS | 413 | 0.158946 | NS |
| WT:PFF-WT:PBS | 62.55556 | 0.999113 | NS |
| Hom:PFF-Het:PFF | 236.4333 | 0.714194 | NS |
| WT:PFF-Het:PFF | -114.011 | 0.983236 | NS |
| WT:PFF-Hom:PFF | -350.444 | 0.320023 | NS |

**Table S22.** Summary statistics (ANOVA and Tukey’s HSD; CI = 95%) for Center Movement.

| Pairwise Comparison | Difference (Tukey) | Adjusted p-value (Tukey) | Significance |
| --- | --- | --- | --- |
| Hom-Het | 316.7 | 0.196382 | NS |
| WT-Het | -17.5778 | 0.994801 | NS |
| WT-Hom | -334.278 | 0.179246 | NS |
| **PFF-PBS** | **410.6071** | **0.008033** | **\*\*** |
| Hom:PBS-Het:PBS | 132.5222 | 0.995205 | NS |
| WT:PBS-Het:PBS | -3.58889 | 1 | NS |
| Het:PFF-Het:PBS | 301.2 | 0.829729 | NS |
| **Hom:PFF-Het:PBS** | **802.0778** | **0.03214** | **\*** |
| WT:PFF-Het:PBS | 269.6333 | 0.896656 | NS |
| WT:PBS-Hom:PBS | -136.111 | 0.995181 | NS |
| Het:PFF-Hom:PBS | 168.6778 | 0.985481 | NS |
| Hom:PFF-Hom:PBS | 669.5556 | 0.128647 | NS |
| WT:PFF-Hom:PBS | 137.1111 | 0.995013 | NS |
| Het:PFF-WT:PBS | 304.7889 | 0.838481 | NS |
| **Hom:PFF-WT:PBS** | **805.6667** | **0.03807** | **\*** |
| WT:PFF-WT:PBS | 273.2222 | 0.901419 | NS |
| Hom:PFF-Het:PFF | 500.8778 | 0.379761 | NS |
| WT:PFF-Het:PFF | -31.5667 | 0.999996 | NS |
| WT:PFF-Hom:PFF | -532.444 | 0.340534 | NS |

**Table S23.** Summary statistics (ANOVA and Tukey’s HSD; CI = 95%) for Center Movement (Male)

| **Pairwise Comparison** | **Difference (Tukey)** | **Adjusted p-value (Tukey)** | **Significance** |
| --- | --- | --- | --- |
| Hom-Het | -127.856 | 0.651514 | NS |
| WT-Het | -160.8 | 0.509797 | NS |
| WT-Hom | -32.9444 | 0.973084 | NS |
| PFF-PBS | -35.5357 | 0.765889 | NS |
| Hom:PBS-Het:PBS | -227.7 | 0.87253 | NS |
| WT:PBS-Het:PBS | -125.144 | 0.989571 | NS |
| Het:PFF-Het:PBS | -76.8 | 0.998811 | NS |
| Hom:PFF-Het:PBS | -104.811 | 0.995407 | NS |
| WT:PFF-Het:PBS | -273.256 | 0.762008 | NS |
| WT:PBS-Hom:PBS | 102.5556 | 0.996324 | NS |
| Het:PFF-Hom:PBS | 150.9 | 0.975934 | NS |
| Hom:PFF-Hom:PBS | 122.8889 | 0.991466 | NS |
| WT:PFF-Hom:PBS | -45.5556 | 0.999929 | NS |
| Het:PFF-WT:PBS | 48.34444 | 0.999891 | NS |
| Hom:PFF-WT:PBS | 20.33333 | 0.999999 | NS |
| WT:PFF-WT:PBS | -148.111 | 0.980191 | NS |
| Hom:PFF-Het:PFF | -28.0111 | 0.999993 | NS |
| WT:PFF-Het:PFF | -196.456 | 0.92745 | NS |
| WT:PFF-Hom:PFF | -168.444 | 0.965371 | NS |

**Table S24.** Summary statistics (ANOVA and Tukey’s HSD; CI = 95%) for Rearing (All).

| Pairwise Comparison | Difference (Tukey) | Adjusted p-value<br>(Tukey) | Significance |
| --- | --- | --- | --- |
| Hom-Het | 0.152778 | 0.999858 | NS |
| WT-Het | -5.01389 | 0.858363 | NS |
| WT-Hom | -5.16667 | 0.857215 | NS |
| PFF-PBS | 8.875 | 0.259481 | NS |
| Hom:PBS-Het:PBS | -3.47778 | 0.999839 | NS |
| WT:PBS-Het:PBS | -3.75556 | 0.999765 | NS |
| Het:PFF-Het:PBS | 7.35 | 0.993269 | NS |
| Hom:PFF-Het:PBS | 11.13333 | 0.961844 | NS |
| WT:PFF-Het:PBS | 1.077778 | 1 | NS |
| WT:PBS-Hom:PBS | -0.27778 | 1 | NS |
| Het:PFF-Hom:PBS | 10.82778 | 0.966117 | NS |
| Hom:PFF-Hom:PBS | 14.61111 | 0.896647 | NS |
| WT:PFF-Hom:PBS | 4.555556 | 0.999468 | NS |
| Het:PFF-WT:PBS | 11.10556 | 0.962247 | NS |
| Hom:PFF-WT:PBS | 14.88889 | 0.888988 | NS |
| WT:PFF-WT:PBS | 4.833333 | 0.99929 | NS |
| Hom:PFF-Het:PFF | 3.783333 | 0.999757 | NS |
| WT:PFF-Het:PFF | -6.27222 | 0.997187 | NS |
| WT:PFF-Hom:PFF | -10.0556 | 0.978035 | NS |

**Table S25.** Summary statistics (ANOVA and Tukey’s HSD; CI = 95%) for Rearing (Female).

| **Pairwise Comparison** | **Difference (Tukey)** | **Adjusted p-value (Tukey)** | **Significance** |
| --- | --- | --- | --- |
| Hom-Het | 7.594444 | 0.84541 | NS |
| WT-Het | -7.68333 | 0.842087 | NS |
| WT-Hom | -15.2778 | 0.528129 | NS |
| PFF-PBS | 19.42857 | 0.091647 | . |
| Hom:PBS-Het:PBS | -4.32222 | 0.99992 | NS |
| WT:PBS-Het:PBS | -8.1 | 0.998289 | NS |
| Het:PFF-Het:PBS | 11.5 | 0.989946 | NS |
| Hom:PFF-Het:PBS | 31.01111 | 0.60447 | NS |
| WT:PFF-Het:PBS | 4.233333 | 0.999928 | NS |
| WT:PBS-Hom:PBS | -3.77778 | 0.999964 | NS |
| Het:PFF-Hom:PBS | 15.82222 | 0.96348 | NS |
| Hom:PFF-Hom:PBS | 35.33333 | 0.491764 | NS |
| WT:PFF-Hom:PBS | 8.555556 | 0.998034 | NS |
| Het:PFF-WT:PBS | 19.6 | 0.91271 | NS |
| Hom:PFF-WT:PBS | 39.11111 | 0.377683 | NS |
| WT:PFF-WT:PBS | 12.33333 | 0.989122 | NS |
| Hom:PFF-Het:PFF | 19.51111 | 0.914236 | NS |
| WT:PFF-Het:PFF | -7.26667 | 0.998985 | NS |
| WT:PFF-Hom:PFF | -26.7778 | 0.759301 | NS |

**Table S26.**
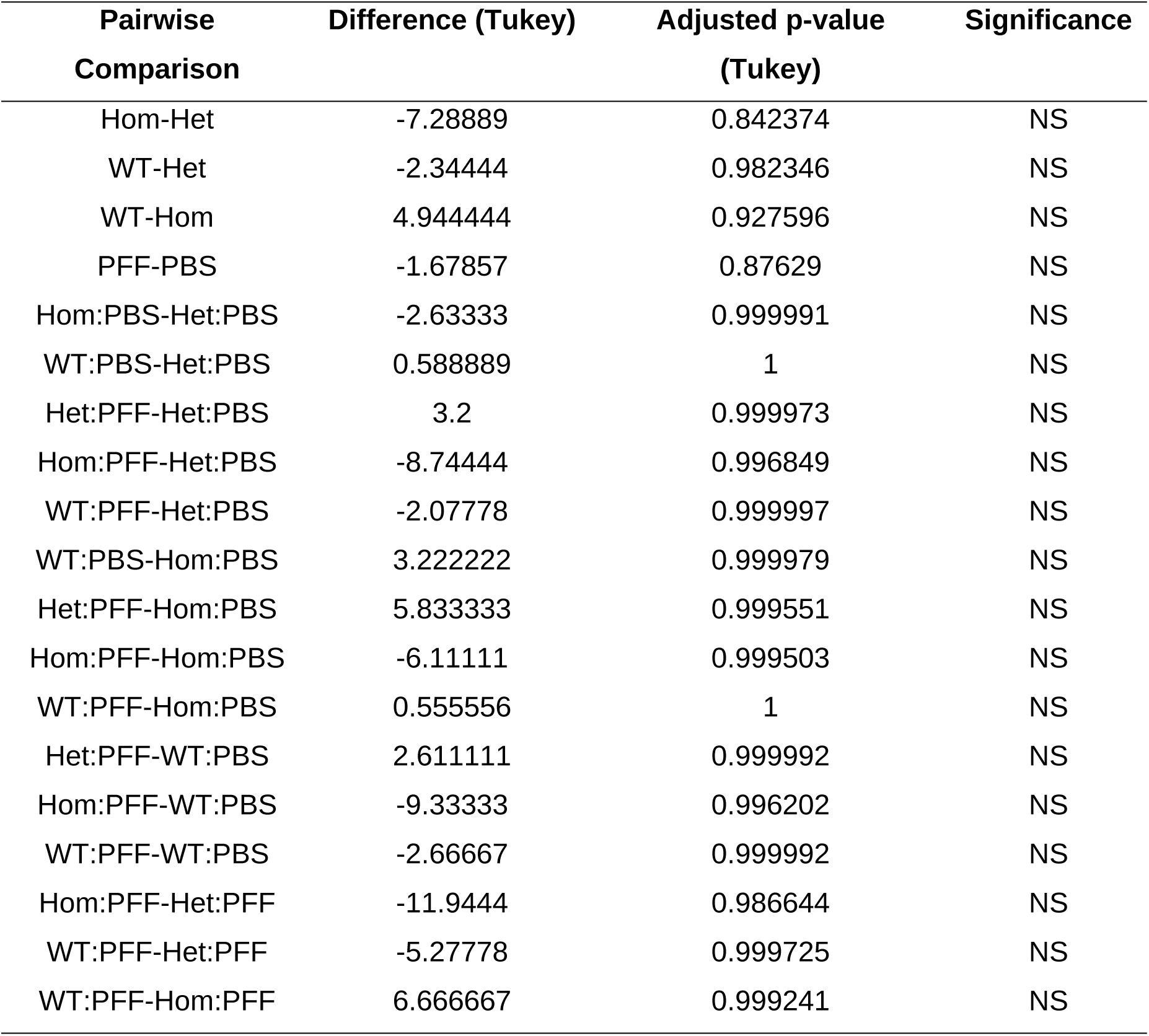
Summary statistics (ANOVA and Tukey’s HSD; CI = 95%) for Rearing (Male).

**Table S27.** Summary statistics (ANOVA and Tukey’s HSD; CI = 95%) for pS129 (IHC - Fluorescence)

| Pairwise Comparison | Difference (Tukey) | Adjusted p-value<br>(Tukey) | Significance |
| --- | --- | --- | --- |
| WT:PBS-Het:PBS | 0.04724 | >0.9999 | NS |
| WT:PBS-Hom:PBS | 0.03456 | >0.9999 | NS |
| **WT:PBS-WT:PFF** | **-11.44** | **<0.0001** | **\*\*\*\*** |
| **WT:PBS-Het:PFF** | **-4.978** | **<0.0001** | **\*\*\*\*** |
| WT:PBS-Hom:PFF | -1.678 | 0.4235 | NS |
| Het:PBS-Hom:PBS | -0.01268 | >0.9999 | NS |
| **Het:PBS-WT:PFF** | **-11.48** | **<0.0001** | **\*\*\*\*** |
| **Het:PBS-Het:PFF** | **-5.025** | **<0.0001** | **\*\*\*\*** |
| Het:PBS-Hom:PFF | -1.726 | 0.3925 | NS |
| **Hom:PBS-WT:PFF** | **-11.47** | **<0.0001** | **\*\*\*\*** |
| **Hom:PBS-Het:PFF** | **-5.012** | **<0.0001** | **\*\*\*\*** |
| Hom:PBS-Hom:PFF | -1.713 | 0.4007 | NS |
| **WT:PFF-Het:PFF** | **6.458** | **<0.0001** | **\*\*\*\*** |
| **WT:PFF-Hom:PFF** | **9.758** | **<0.0001** | **\*\*\*\*** |
| **Het:PFF-Hom:PFF** | **3.299** | **0.0071** | **\*\*** |

**Table S28.** Summary statistics (ANOVA and Tukey’s HSD; CI = 95%) for Th(IHC-Fluorescence)

| Pairwise Comparison | Difference (Tukey) | Adjusted p-value<br>(Tukey) | Significance |
| --- | --- | --- | --- |
| WT:PBS-Het:PBS | 0.2822 | 0.9994 | NS |
| WT:PBS-Hom:PBS | 0.4757 | 0.9924 | NS |
| **WT:PBS-WT:PFF** | **19.16** | **<0.0001** | **\*\*\*\*** |
| **WT:PBS-Het:PFF** | **14.66** | **<0.0001** | **\*\*\*\*** |
| **WT:PBS-Hom:PFF** | **7.534** | **<0.0001** | **\*\*\*\*** |
| Het:PBS-Hom:PBS | 0.1934 | >0.9999 | NS |
| **Het:PBS-WT:PFF** | **18.88** | **<0.0001** | **\*\*\*\*** |
| **Het:PBS-Het:PFF** | **14.37** | **<0.0001** | **\*\*\*\*** |
| **Het:PBS-Hom:PFF** | **7.252** | **<0.0001** | **\*\*\*\*** |
| **Hom:PBS-WT:PFF** | **18.68** | **<0.0001** | **\*\*\*\*** |
| **Hom:PBS-Het:PFF** | **14.18** | **<0.0001** | **\*\*\*\*** |
| **Hom:PBS-Hom:PFF** | **7.058** | **<0.0001** | **\*\*\*\*** |
| **WT:PFF-Het:PFF** | **-4.502** | **<0.0001** | **\*\*\*\*** |
| **WT:PFF-Hom:PFF** | **-11.62** | **<0.0001** | **\*\*\*\*** |
| **Het:PFF-Hom:PFF** | **-7.122** | **<0.0001** | **\*\*\*\*** |

**Table S29.** Summary statistics (ANOVA and Tukey’s HSD; CI = 95%) for pS129/Th ratio (IHC-Fluorescence)

| Pairwise Comparison | Difference (Tukey) | Adjusted p-value<br>(Tukey) | Significance |
| --- | --- | --- | --- |
| WT:PBS-Het:PBS | 0.001333 | >0.9999 | NS |
| WT:PBS-Hom:PBS | 0.0001193 | >0.9999 | NS |
| **WT:PBS-WT:PFF** | **-3.071** | **<0.0001** | **\*\*\*\*** |
| WT:PBS-Het:PFF | -0.6681 | 0.1382 | NS |
| WT:PBS-Hom:PFF | -0.1368 | 0.9952 | NS |
| Het:PBS-Hom:PBS | -0.001213 | >0.9999 | NS |
| **Het:PBS-WT:PFF** | **-3.072** | **<0.0001** | **\*\*\*\*** |
| Het:PBS-Het:PFF | -0.6695 | 0.1368 | NS |
| Het:PBS-Hom:PFF | -0.1381 | 0.9950 | NS |
| **Hom:PBS-WT:PFF** | **-3.071** | **<0.0001** | **\*\*\*\*** |
| Hom:PBS-Het:PFF | -0.6683 | 0.1381 | NS |
| Hom:PBS-Hom:PFF | -0.1369 | 0.9952 | NS |
| **WT:PFF-Het:PFF** | **2.403** | **<0.0001** | **\*\*\*\*** |
| **WT:PFF-Hom:PFF** | **2.934** | **<0.0001** | **\*\*\*\*** |
| Het:PFF-Hom-PFF | 0.5313 | 0.3536 | NS |

**Table S30.** Summary statistics (ANOVA and Tukey’s HSD; CI = 95%) for Th (IHC)

| **Pairwise**<br>**Comparison** | **Difference (Tukey)** | **Adjusted p-value**<br>**(Tukey)** | **Significance** |
| --- | --- | --- | --- |
| WT:PBS-Het:PBS | 28.73 | >0.9999 | NS |
| WT:PBS-Hom:PBS | -54.97 | >0.9999 | NS |
| **WT:PBS-WT:PFF** | **4685** | **<0.0001** | **\*\*\*\*** |
| **WT:PBS-Het:PFF** | **3138** | **<0.0001** | **\*\*\*\*** |
| **WT:PBS-Hom:PFF** | **1924** | **<0.0001** | **\*\*\*\*** |
| Het:PBS-Hom:PBS | -83.70 | 0.9999 | NS |
| **Het:PBS-WT:PFF** | **4657** | **<0.0001** | **\*\*\*\*** |
| **Het:PBS-Het:PFF** | **3109** | **<0.0001** | **\*\*\*\*** |
| **Het:PBS-Hom:PFF** | **1895** | **<0.0001** | **\*\*\*\*** |
| **Hom:PBS-WT:PFF** | **4740** | **<0.0001** | **\*\*\*\*** |
| **Hom:PBS-Het:PFF** | **3193** | **<0.0001** | **\*\*\*\*** |
| **Hom:PBS-Hom:PFF** | **1979** | **<0.0001** | **\*\*\*\*** |
| **WT:PFF-Het:PFF** | **-1547** | **0.0009** | **\*\*\*** |
| **WT:PFF-Hom:PFF** | **-2761** | **<0.0001** | **\*\*\*\*** |
| **Het:PFF-Hom:PFF** | **-1214** | **0.0152** | **\*** |

**Table S31.** Summary statistics (ANOVA and Tukey’s HSD; CI = 95%) for Nissl (IHC)

| **Pairwise Comparison** | **Difference**<br>**(Tukey)** | **Adjusted p-value**<br>**(Tukey)** | **Significance** |
| --- | --- | --- | --- |
| WT-PBS-Het:PBS | -11.08 | >0.9999 | NS |
| WT:PBS-Hom:PBS | -75.88 | >0.9999 | NS |
| **WT:PBS-WT:PFF** | **5108** | **<0.0001** | **\*\*\*\*** |
| **WT:PBS-Het:PFF** | **3396** | **<0.0001** | **\*\*\*\*** |
| **WT:PBS-Hom:PFF** | **2061** | **<0.0001** | **\*\*\*\*** |
| Het:PBS-Hom:PBS | -64.80 | >0.9999 | NS |
| **Het:PBS-WT:PFF** | **5119** | **<0.0001** | **\*\*\*\*** |
| **Het:PBS-Het:PFF** | **3407** | **<0.0001** | **\*\*\*\*** |
| **Het:PBS-Hom:PFF** | **2072** | **<0.0001** | **\*\*\*\*** |
| **Hom:PBS-WT:PFF** | **5184** | **<0.0001** | **\*\*\*\*** |
| **Hom:PBS-Het:PFF** | **3472** | **<0.0001** | **\*\*\*\*** |
| **Hom:PBS-Hom:PFF** | **2137** | **<0.0001** | **\*\*\*\*** |
| WT:PFF-Het:PFF | -1712 | 0.0008 | *** |
| **WT:PFF-Hom:PFF** | **-3047** | **<0.0001** | **\*\*\*\*** |
| **Het:PFF-Hom:PFF** | **-1335** | **0.0143** | **\*** |

**Table S32.** Summary statistics (ANOVA and Tukey’ s HSD; CI = 95%) for Th (WB)

| Pairwise Comparison | Difference (Tukey) | Adjusted p-value<br>(Tukey) | Significance |
| --- | --- | --- | --- |
| WT:PBS-Het:PBS | -0.03292 | 0.9996 | NS |
| WT:PBS-Hom:PBS | 0.1174 | 0.8785 | NS |
| **WT:PBS-WT:PFF** | **1.035** | **<0.0001** | **\*\*\*\*** |
| **WT:PBS-Het:PFF** | **0.7225** | **<0.0001** | **\*\*\*\*** |
| **WT:PBS-Hom:PFF** | **0.4031** | **0.0083** | **\*\*** |
| Het:PBS-Hom:PBS | 0.1503 | 0.7235 | NS |
| **Het:PBS-WT:PFF** | **1.068** | **<0.0001** | **\*\*\*\*** |
| **Het:PBS-Het:PFF** | **0.7554** | **<0.0001** | **\*\*\*\*** |
| **Het:PBS-Hom:PFF** | **0.4360** | **0.0037** | **\*\*** |
| **Hom:PBS-WT:PFF** | **0.9176** | **<0.0001** | **\*\*\*\*** |
| **Hom:PBS-Het:PFF** | **0.6051** | **<0.0001** | **\*\*\*\*** |
| Hom:PBS-Hom:PFF | 0.2857 | 0.1105 | NS |
| **WT:PFF-Het:PFF** | **-0.3125** | **0.0408** | **\*** |
| **WT:PFF-Hom:PFF** | **-0.6319** | **<0.0001** | **\*\*\*\*** |
| Het:PFF-Hom:PFF | -0.3194 | 0.0501 | . |

**Table S33.**
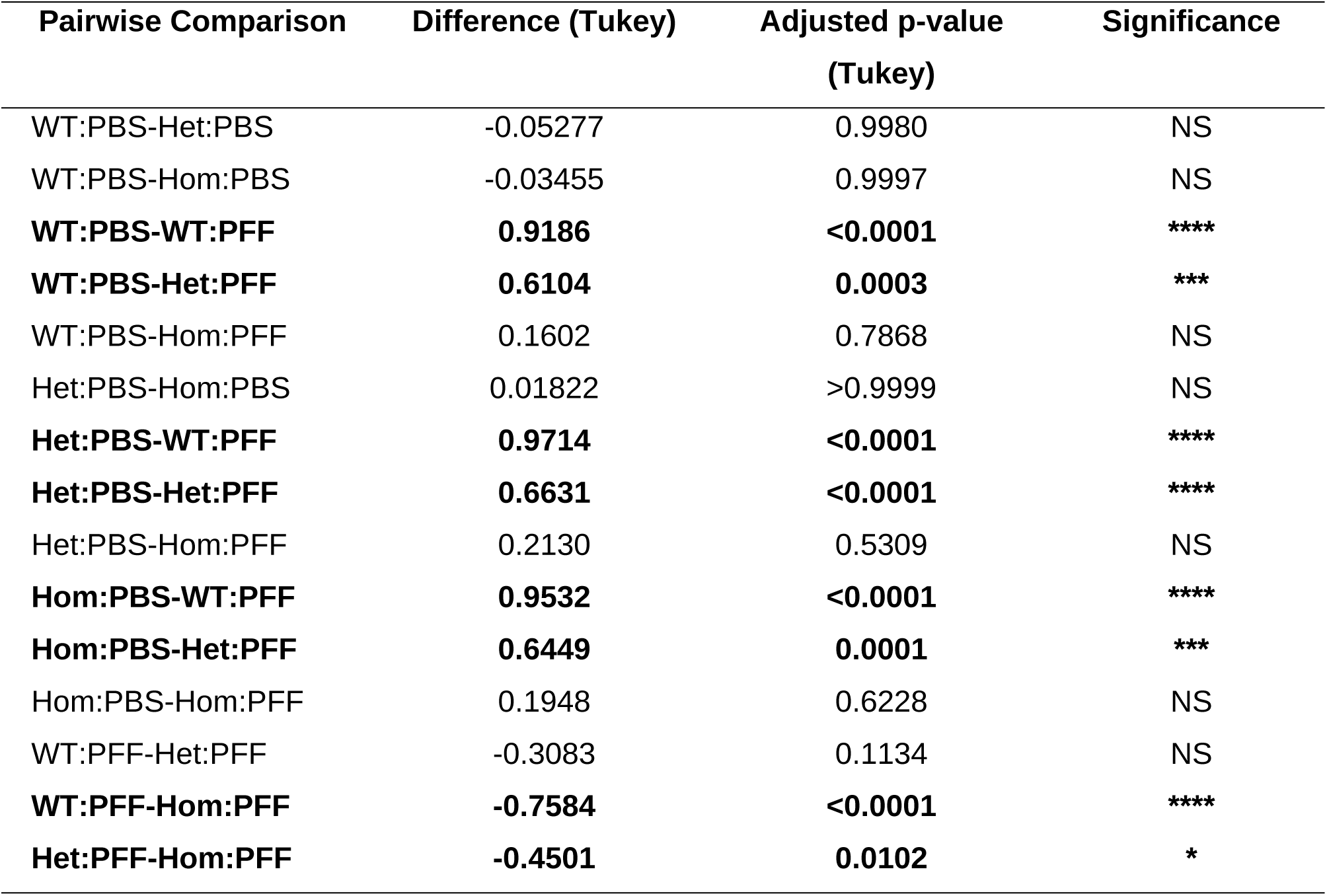
Summary statistics (ANOVA and Tukey’s HSD; CI = 95%) for DAT (WB)

**Table S34.** Summary statistics (ANOVA and Tukey’s HSD; CI = 95%) for insoluble α-Syn (WB)

| Pairwise Comparison | Difference (Tukey) | Adjusted p-value<br>(Tukey) | Significance |
| --- | --- | --- | --- |
| WT:PBS-Het:PBS | 0.05434 | >0.9999 | NS |
| WT:PBS-Hom:PBS | -0.02308 | >0.9999 | NS |
| **WT:PBS-WT:PFF** | **-2.730** | **<0.0001** | **\*\*\*\*** |
| **WT:PBS-Het:PFF** | **-1.398** | **0.0003** | **\*\*\*** |
| WT:PBS-Hom:PFF | -0.4173 | 0.6821 | NS |
| Het:PBS-Hom:PBS | -0.07742 | 0.9998 | NS |
| **Het:PBS-WT:PFF** | **-2.785** | **<0.0001** | **\*\*\*\*** |
| **Het:PBS-Het:PFF** | **-1.452** | **0.0002** | **\*\*\*** |
| Het:PBS-Hom:PFF | -0.4717 | 0.5627 | NS |
| **Hom:PBS-WT:PFF** | **-2.707** | **<0.0001** | **\*\*\*\*** |
| **Hom:PBS-Het:PFF** | **-1.375** | **0.0003** | **\*\*\*** |
| Hom:PBS-Hom:PFF | -0.3943 | 0.7306 | NS |
| **WT:PFF-Het:PFF** | **1.332** | **0.0003** | **\*\*\*** |
| **WT:PFF-Hom:PFF** | **2.313** | **<0.0001** | **\*\*\*\*** |
| **Het:PFF-Hom:PFF** | **0.9808** | **0.0155** | **\*** |

**Table S35.** Summary statistics (ANOVA and Tukey’s HSD; CI = 95%) for insoluble pS129-α-Syn (WB)

| Pairwise Comparison | Difference (Tukey) | Adjusted p-value<br>(Tukey) | Significance |
| --- | --- | --- | --- |
| WT:PBS-Het:PBS | 0.04770 | >0.9999 | NS |
| WT:PBS-Hom:PBS | -0.03628 | >0.9999 | NS |
| **WT:PBS-WT:PFF** | **-3.120** | **<0.0001** | **\*\*\*\*** |
| **WT:PBS-Het:PFF** | **-1.277** | **0.0020** | **\*\*** |
| WT:PBS-Hom:PFF | -0.3925 | 0.7817 | NS |
| Het:PBS-Hom:PBS | -0.08399 | 0.9997 | NS |
| **Het:PBS-WT:PFF** | **-3.168** | **<0.0001** | **\*\*\*\*** |
| **Het:PBS-Het:PFF** | **-1.325** | **0.0013** | **\*\*** |
| Het:PBS-Hom:PFF | -0.4402 | 0.6910 | NS |
| **Hom:PBS-WT:PFF** | **-3.084** | **<0.0001** | **\*\*\*\*** |
| **Hom:PBS-Het:PFF** | **-1.241** | **0.0028** | **\*\*** |
| Hom:PBS-Hom:PFF | -0.3562 | 0.8421 | NS |
| **WT:PFF-Het:PFF** | **1.843** | **<0.0001** | **\*\*\*\*** |
| **WT:PFF-Hom:PFF** | **2.728** | **<0.0001** | **\*\*\*\*** |
| Het:PFF-Hom:PFF | 0.8844 | 0.0566 | NS |

**Table S36.**
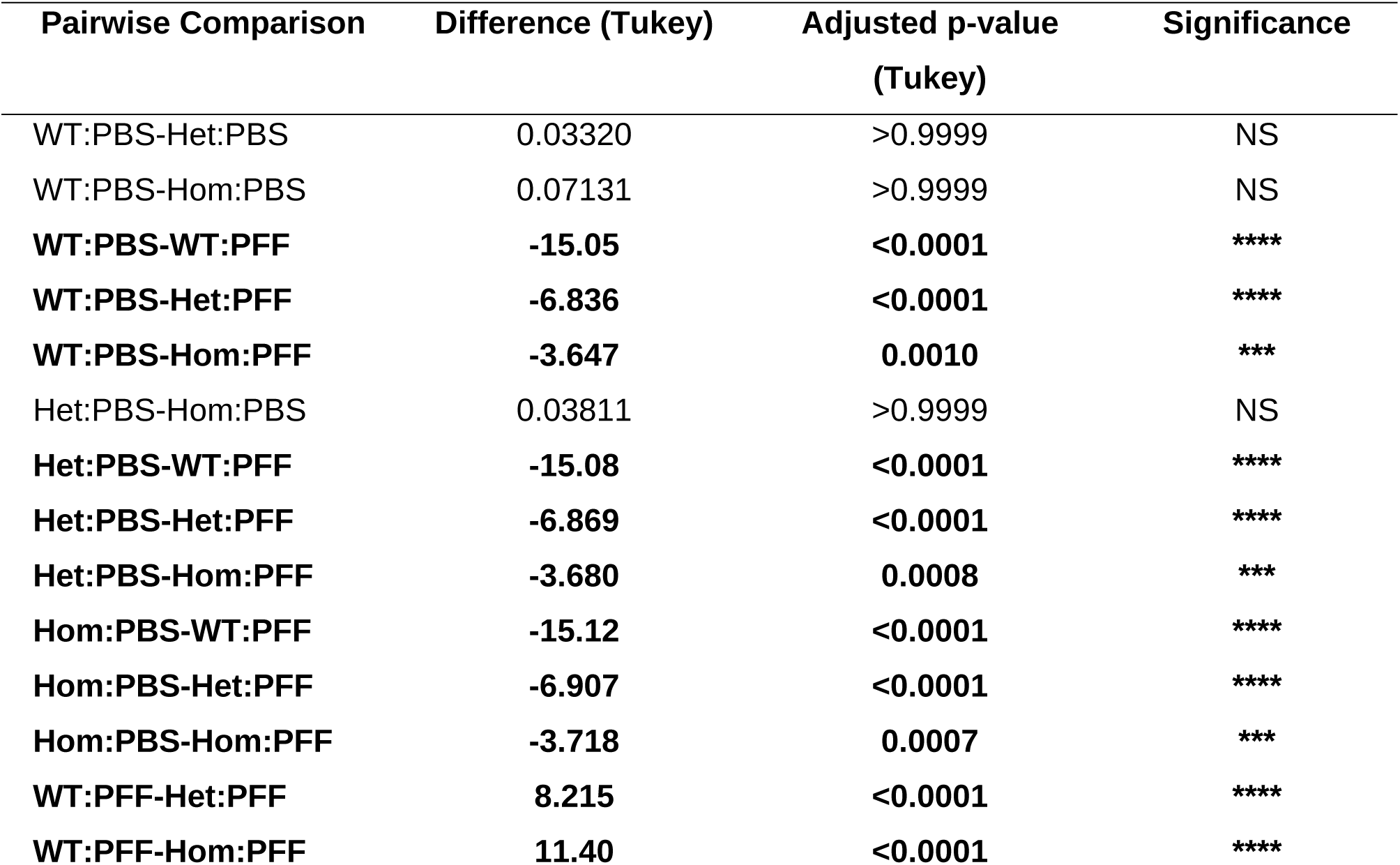
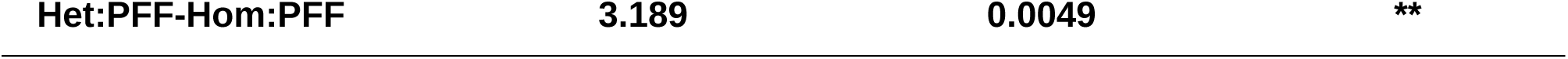
Summary statistics (ANOVA and Tukey’s HSD; CI = 95%) for Iba1 (IHC-Fluorescence)

**Table S37.**
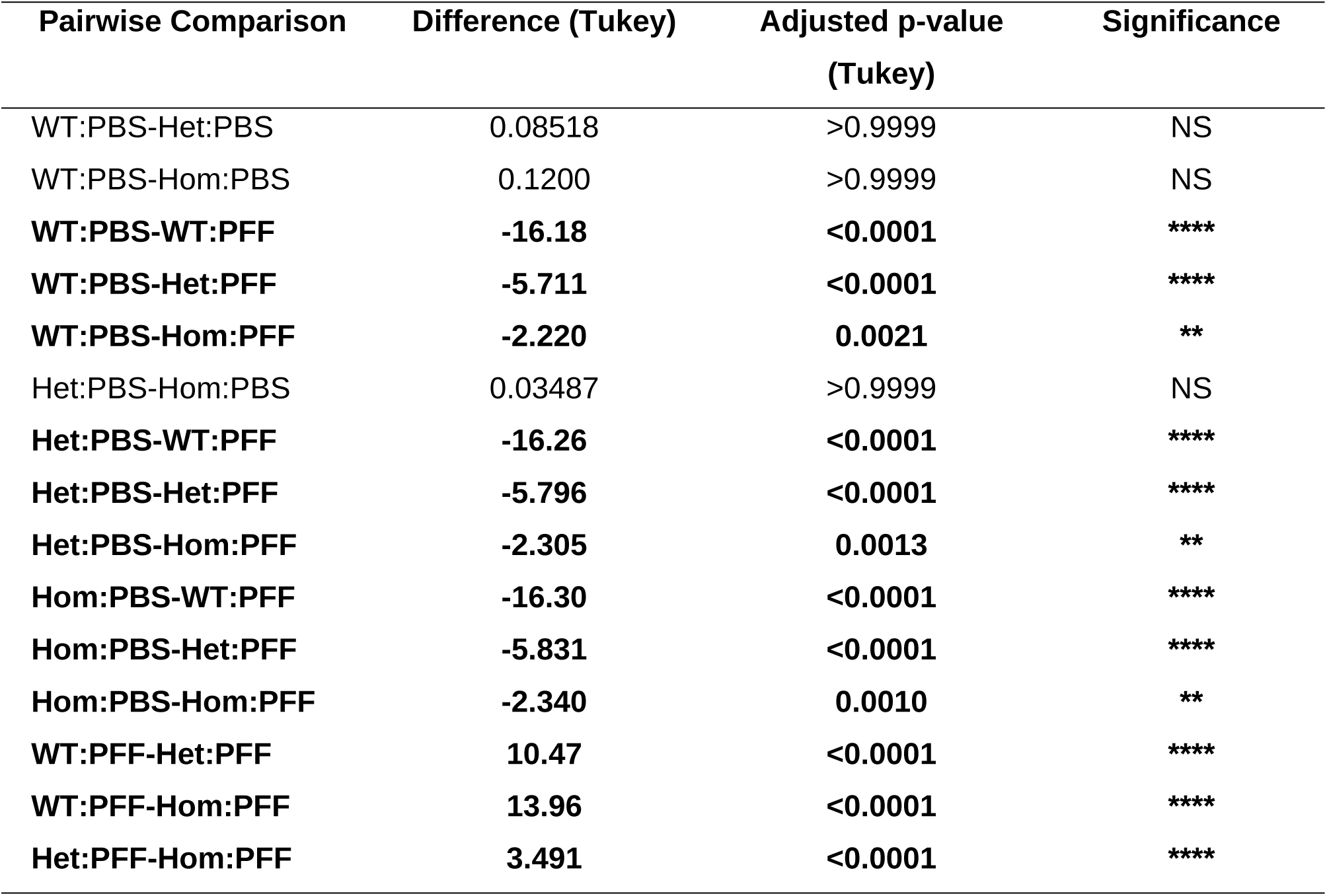
Summary statistics (ANOVA and Tukey’s HSD; CI = 95%) for Gfap (IHC-Fluorescence)

**Table S38.** Summary statistics (ANOVA and Tukey’s HSD; CI = 95%) for Iba1 (WB)

| Pairwise Comparison | Difference (Tukey) | Adjusted p-value<br>(Tukey) | Significance |
| --- | --- | --- | --- |
| WT:PBS-Het:PBS | -0.07133 | 0.9995 | NS |
| WT:PBS-Hom:PBS | -0.1448 | 0.9858 | NS |
| **WT:PBS-WT:PFF** | **-2.738** | **<0.0001** | **\*\*\*\*** |
| **WT:PBS-Het:PFF** | **-1.704** | **<0.0001** | **\*\*\*\*** |
| WT:PBS-Hom:PFF | -0.4281 | 0.4056 | NS |
| Het:PBS-Hom:PBS | -0.07345 | 0.9994 | NS |
| **Het:PBS-WT:PFF** | **-2.667** | **<0.0001** | **\*\*\*\*** |
| **Het:PBS-Het:PFF** | **-1.633** | **<0.0001** | **\*\*\*\*** |
| Het:PBS-Hom:PFF | -0.3568 | 0.6009 | NS |
| **Hom:PBS-WT:PFF** | **-2.594** | **<0.0001** | **\*\*\*\*** |
| **Hom:PBS-Het:PFF** | **-1.560** | **<0.0001** | **\*\*\*\*** |
| Hom:PBS-Hom:PFF | -0.2833 | 0.7958 | NS |
| **WT:PFF-Het:PFF** | **1.034** | **0.0003** | **\*\*\*** |
| **WT:PFF-Hom:PFF** | **2.310** | **<0.0001** | **\*\*\*\*** |
| **Het:PFF-Hom:PFF** | **1.276** | **<0.0001** | **\*\*\*\*** |

**Table S39.** Summary statistics (ANOVA and Tukey’s HSD; CI = 95%) for Gfap (WB)

| Pairwise Comparison | Difference (Tukey) | Adjusted p-value<br>(Tukey) | Significance |
| --- | --- | --- | --- |
| WT:PBS-Het:PBS | -0.01154 | >0.9999 | NS |
| WT:PBS-Hom:PBS | -0.07291 | >0.9999 | NS |
| **WT:PBS-WT:PFF** | **-3.205** | **<0.0001** | **\*\*\*\*** |
| WT:PBS-Het:PFF | -1.105 | 0.2889 | NS |
| WT:PBS-Hom:PFF | -0.2693 | 0.9951 | NS |
| Het:PBS-Hom:PBS | -0.06137 | >0.9999 | NS |
| **Het:PBS-WT:PFF** | **-3.194** | **<0.0001** | **\*\*\*\*** |
| Het:PBS-Het:PFF | -1.094 | 0.2995 | NS |
| Het:PBS-Hom:PFF | -0.2578 | 0.9960 | NS |
| **Hom:PBS-WT:PFF** | **-3.132** | **<0.0001** | **\*\*\*\*** |
| Hom:PBS-Het:PFF | -1.032 | 0.3602 | NS |
| Hom:PBS-Hom:PFF | -0.1964 | 0.9989 | NS |
| **WT:PFF-Het:PFF** | **2.100** | **0.0020** | **\*\*** |
| **WT:PFF-Hom:PFF** | **2.936** | **<0.0001** | **\*\*\*\*** |
| Het:PFF-Hom:PFF | 0.8360 | 0.5877 | NS |

## REFERENCES

1. Dorsey, E. R. & Bloem, B. R. The Parkinson pandemic - A call to action. JAMA Neurol. 75, 9–10 (2018).

2. Nalls, M. A. et al. Identification of novel risk loci, causal insights, and heritable risk for Parkinson’s disease: a meta-analysis of genome-wide association studies. Lancet Neurol. 18, 1091–1102 (2019).

3. Fearnley, J. M. & Lees, A. J. Ageing and Parkinson’s disease: Substantia nigra regional selectivity. Brain 114, 2283–2301 (1991).

4. Goedert, M., Spillantini, M. G., Tredici, K. D. & Braak, H. 100 years of Lewy pathology. Nat. Rev. Neurol. 9, 13–24 (2012).

5. Butler, B., Sambo, D. & Khoshbouei, H. Alpha-synuclein modulates dopamine neurotransmission. J. Chem. Neuroanat. 83–84, 41–49 (2017).

6. Gao, V., Briano, J. A., Komer, L. E. & Burré, J. Functional and Pathological Effects of α-Synuclein on Synaptic SNARE Complexes. J. Mol. Biol. 435, 167714 (2023).

7. Stefanis, L. α-Synuclein in Parkinson’s Disease. Cold Spring Harb. Perspect. Med. 2, (2012).

8. Ibáñez, P. et al. Causal relation between alpha-synuclein gene duplication and familial Parkinson’s disease. The Lancet 364, 1169–1171 (2004).

9. Gao, H.-M. et al. Neuroinflammation and α-synuclein dysfunction potentiate each other, driving chronic progression of neurodegeneration in a mouse model of Parkinson’s disease. Environ. Health Perspect. 119, 807 (2011).

10. Polymeropoulos, M. H. et al. Mutation in the α-synuclein gene identified in families with Parkinson’s disease. Science 276, 2045–2047 (1997).

11. Hirsch, E. C., Vyas, S. & Hunot, S. Neuroinflammation in Parkinson’s disease. Parkinsonism Relat. Disord. 18, S210–S212 (2012).

12. Boyd, R. J., Avramopoulos, D., Jantzie, L. L. & McCallion, A. S. Neuroinflammation represents a common theme amongst genetic and environmental risk factors for Alzheimer and Parkinson diseases. J. Neuroinflammation 19, (2022).

13. Nalls, M. A. et al. Large-scale meta-analysis of genome-wide association data identifies six new risk loci for Parkinson’s disease. Nat. Genet. 46, 989–993 (2014).

14. Kim, J. J. et al. Multi-ancestry genome-wide association meta-analysis of Parkinson’s disease. Nat. Genet. 56, 27–36 (2024).

15. Maurano, M. T. et al. Systematic localization of common disease-associated variation in regulatory DNA. Science 337, 1190–1195 (2012).

16. Soldner, F. et al. Parkinson-associated risk variant in distal enhancer of α-synuclein modulates target gene expression. Nature 533, 95–99 (2016).

17. McClymont, S. A. et al. Parkinson-associated SNCA enhancer variants revealed by open chromatin in mouse dopamine neurons. Am. J. Hum. Genet. 103, 874–892 (2018).

18. Daubner, S. C., Le, T. & Wang, S. Tyrosine Hydroxylase and Regulation of Dopamine Synthesis. Arch. Biochem. Biophys. 508, 1 (2011).

19. Volpicelli-Daley, L. A., Luk, K. C. & Lee, V. M. Y. Addition of exogenous α-synuclein preformed fibrils to primary neuronal cultures to seed recruitment of endogenous α-synuclein to Lewy body and Lewy neurite-like aggregates. Nat. Protoc. 9, 2135–2146 (2014).

20. Kim, S. et al. Transneuronal Propagation of Pathologic α-Synuclein from the Gut to the Brain Models Parkinson’s disease. Neuron 103, 627 (2019).

21. Yun, S. P. et al. Block of A1 astrocyte conversion by microglia is neuroprotective in models of Parkinson’s disease. Nat. Med. 24, 931 (2018).

22. Verma, D. K. et al. Alpha-synuclein preformed fibrils induce cellular senescence in parkinson’s disease models. Cells 10, (2021).

23. Mao, X. et al. Pathological α-synuclein transmission initiated by binding lymphocyte-activation gene 3. Science 353, (2016).

24. Ma, S.-X. et al. Complement and Coagulation Cascades are Potentially Involved in Dopaminergic Neurodegeneration in α-Synuclein-Based Mouse Models of Parkinson’s Disease. J. Proteome Res. 20, 3443 (2021).

25. Lee, S. et al. The c-Abl inhibitor, Radotinib HCl, is neuroprotective in a preclinical Parkinson’s disease mouse model. Hum. Mol. Genet. 27, 2344 (2018).

26. Brahmachari, S. et al. Parkin interacting substrate zinc finger protein 746 is a pathological mediator in Parkinson’s disease. Brain 142, 2380 (2019).

27. Luk, K. C. et al. Pathological α-synuclein transmission initiates Parkinson-like neurodegeneration in nontransgenic mice. Science 338, 949–953 (2012).

28. Taylor, T. N., Greene, J. G. & Miller, G. W. Behavioral phenotyping of mouse models of Parkinson’s Disease. Behav. Brain Res. 211, 1–10 (2010).

29. Mahul-Mellier, A.-L. et al. The process of Lewy body formation, rather than simply α-synuclein fibrillization, is one of the major drivers of neurodegeneration. Proc. Natl. Acad. Sci. 117, 4971–4982 (2020).

30. Postuma, R. B. & Berg, D. Advances in markers of prodromal Parkinson disease. Nat. Rev. Neurol. 12, 622–634 (2016).

## REFERENCES

1. Watkins-Chow, D. E. et al. Highly Efficient Cpf1-Mediated Gene Targeting in Mice Following High Concentration Pronuclear Injection. G3 (Bethesda, Md.) 7, 719–722 (2017).

2. Taylor, S. C. et al. The ultimate qPCR experiment: Producing publication quality, reproducible data the first time. Trends in Biotechnology 37, 761–774 (2019).

3. Volpicelli-Daley, L. A., Luk, K. C. & Lee, V. M. Y. Addition of exogenous α-Synuclein Pre-formed fibrils to Primary Neuronal Cultures to seed recruitment of endogenous α-Synuclein to Lewy body and Lewy Neurite-like aggregates. Nature protocols 9, 2135 (2014).

4. Kim, S. et al. Transneuronal Propagation of Pathologic α-Synuclein from the Gut to the Brain Models Parkinson’s disease. Neuron 103, 627 (2019).

5. Yun, S. P. et al. Block of A1 astrocyte conversion by microglia is neuroprotective in models of Parkinson’s disease. Nature medicine 24, 931 (2018).

6. Verma, D. K. et al. Alpha-synuclein preformed fibrils induce cellular senescence in parkinson’s disease models. Cells 10, (2021).

7. Mao, X. et al. Pathological α-synuclein transmission initiated by binding lymphocyte-activation gene 3. Science 353, (2016).

8. Lee, S. et al. The c-Abl inhibitor, Radotinib HCl, is neuroprotective in a preclinical Parkinson’s disease mouse model. Human Molecular Genetics 27, 2344 (2018).

9. Kim, D. et al. Graphene quantum dots prevent α-synucleinopathy in Parkinson’s disease. Nature nanotechnology 13, 812 (2018).

10. Kam, T. I. et al. Poly (ADP-ribose) Drives Pathologic α-Synuclein Neurodegeneration in Parkinson’s Disease. Science 362, (2018).

11. Lee, Y. et al. Parthanatos Mediates AIMP2 Activated Age Dependent Dopaminergic Neuronal Loss. Nature neuroscience 16, 1392 (2013).

